# A conserved transcription factor controls gluconeogenesis via distinct targets in hypersaline-adapted archaea with diverse metabolic capabilities

**DOI:** 10.1101/2023.08.19.553955

**Authors:** Rylee K. Hackley, Angie Vreugdenhil-Hayslette, Cynthia L. Darnell, Amy K. Schmid

## Abstract

Timely regulation of carbon metabolic pathways is essential for cellular processes and to prevent futile cycling of intracellular metabolites. In *Halobacterium salinarum*, a hypersaline adapted archaeon, a sugar-sensing TrmB family protein controls gluconeogenesis and other biosynthetic pathways. Notably, *Hbt. salinarum* does not utilize carbohydrates for energy, uncommon among Haloarchaea. We characterized a TrmB-family transcriptional regulator in a saccharolytic generalist, *Haloarcula hispanica*, to investigate whether the targets and function of TrmB, or its regulon, is conserved in related species with distinct metabolic capabilities. In *Har. hispanica*, TrmB binds to 15 sites across the genome and induces the expression of genes primarily involved in gluconeogenesis and tryptophan biosynthesis. An important regulatory control point in *Hbt. salinarum*, activation of *ppsA* and repression of *pykA*, is absent in *Har. hispanica*. Contrary to its role in *Hbt. salinarum* and saccharolytic hyperthermophiles, TrmB does not act as a global regulator: it does not directly repress the expression of glycolytic enzymes, peripheral pathways such as cofactor biosynthesis, or catabolism of other carbon sources in *Har. hispanica*. Cumulatively, these findings suggest re-wiring of the TrmB regulon alongside metabolic network evolution in Haloarchaea.

## Introduction

Regulation of glycolytic and gluconeogenic activities in the cell is critical to generate energy and direct carbon flux in the face of variable environments and nutrient availability. In bacteria and eukaryotes, allosteric regulation plays an important role, though regulation at the transcriptional and post-transcriptional levels also occurs [1–4]. Studies in archaea, however, suggest that allosteric regulation of enzymes involved in central carbon metabolism is less prevalent (reviewed in Ref. [5]). For example, reactions catalyzed by the antagonistic enzyme couples phosphofructokinase and fructose-1,6-bisphosphatase are not allosterically regulated in most characterized archaeal enzymes, as they are in bacteria. Archaeal pyruvate kinases appear to be sensitive to allosteric activation by novel ligands AMP and 3-phosphoglycerate [6–8]. Instead, studies comparing glycolytic and gluconeogenic growth conditions in archaea indicate regulation at the transcriptional level to be important [9–16].

In *Pyrococcus furiosus* and *Themococcus kodakarensis*, both hyperthermophilic members of Euryarchaea, a conserved transcription factor (TF) controls gluconeogenesis. TrmBL1 in *Pyr. furiosus* and Tgr in *Tcc. kodakarensis* are TrmB family regulators that bind DNA in the absence of glucose to induce the expression of gluconeogenic genes and suppress the expression of glycolytic genes. Both homologs recognize a conserved cis-regulatory motif that is absent from closely related, non-carbohydrate utilizing species [17–19]. The direction of regulation is determined by the motif location relative to other promoter elements: binding downstream of the TATA-box inhibits RNA polymerase recruitment, whereas upstream binding activates transcription.

Though this class of sugar-sensing TrmB regulators is broadly conserved in archaeal and bacterial lineages [20, 21], the majority of homologs are found in Halobacteria, a class of hypersaline-adapted Euryarchaea (Fig 1A) [22]. The non-carbohydrate utilizing, or nonsaccharolytic, species *Halobacterium salinarum NRC-1* encodes a single TrmB homolog that has been characterized previously. In *Hbt. salinarum*, TrmB regulates more than 100 genes via the same mode of regulation proposed for hyperthermophiles. TrmB regulates the expression of genes in the absence of glucose to activate gluconeogenesis and suppress other metabolic pathways such as amino acid metabolism, cobalamin biosynthesis, and purine biosynthesis [10, 23]. Although glucose is not actively transported into the cell nor catabolized via glycolysis, it is essential for glycosylation of the proteinaceous cell surface layer (s-layer) [24–28]. TrmB plays an essential role in this process by controlling the availability of glucose moieties and expression of enzymes for amino acid metabolism and co-factor biosynthesis, and decreasing flux through purine biosynthetic pathways [10, 23, 29, 30]. The genes encoding phosphoenolpyruvate synthase and pyruvate kinase comprise an important regulatory point controlling carbon flux in *Hbt. salinarum*. Within five minutes of glucose addition, *ppsA* is de-activated and *pykA* is de-repressed by TrmB [23]. Furthermore, TrmB regulates the expression of several other transcription factors, endowing the gene regulatory network (GRN) with dynamical properties such as transient ”just-in-time” expression of metabolic genes during rapid nutrient shifts, suggesting that TrmB is a hub at the core of a larger GRN.

**Fig 1.**
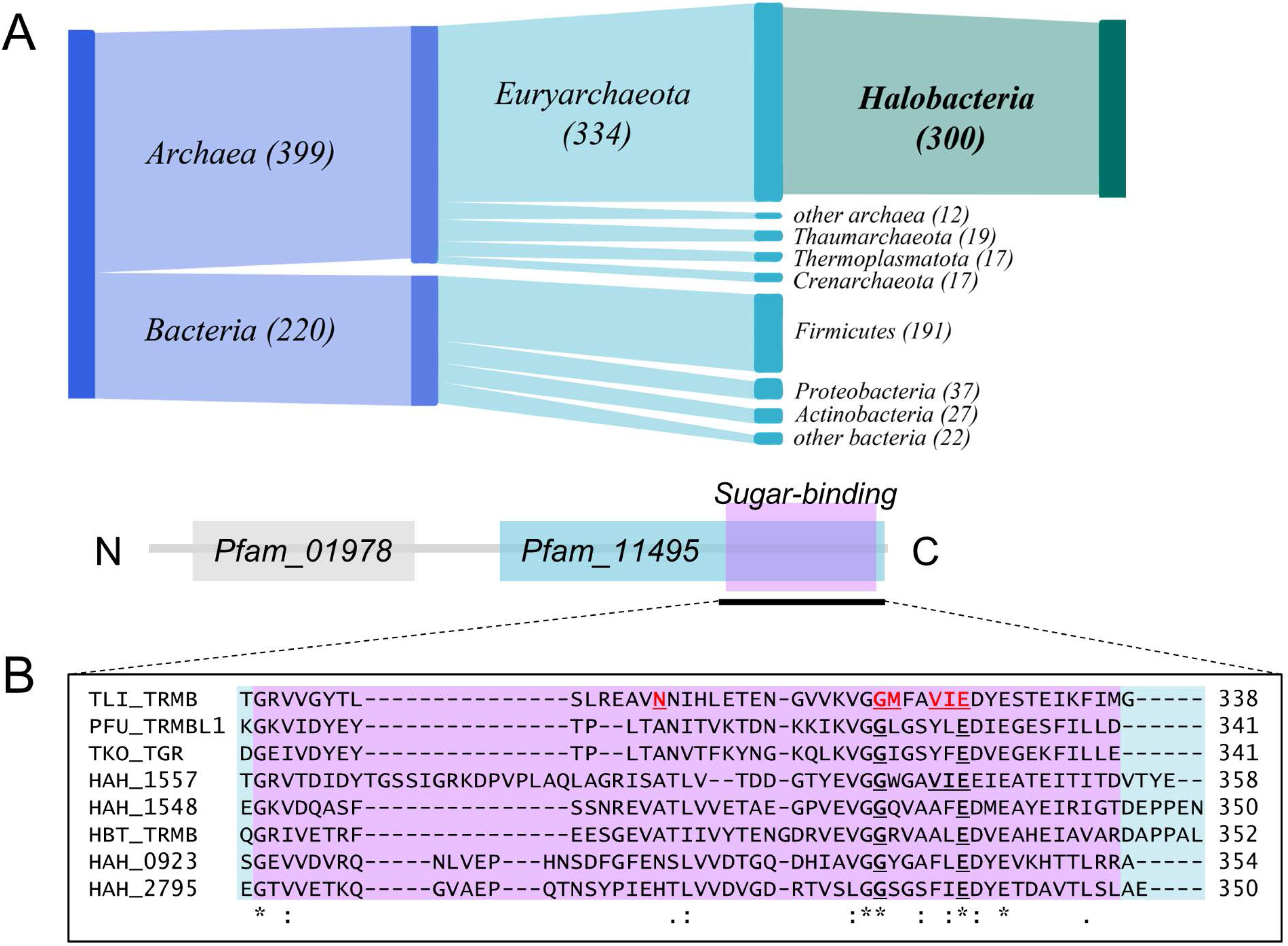
Sugar-sensing TrmB homologs. A) Phylogenetic distribution of TrmB proteins that also contain a carbohydrate-binding domain. B) Multiple sequence alignment of the sugar-binding domain of *Har. hispanica* TrmB homologs. Red, underlined residues are essential for sugar binding in *Tcc. litoralis*, conserved residues are in bold underline [72, 73]. Asterisks below the alignment denote identical residues and dots represent similar residues. Organism abbreviations and locus tags are as follows: TLI TRMB, *Tcc. litoralis* OCC 03542; PFU TRMBL1, *Pyr. furiosus* PF0124; TKO TGR, *Tcc. kodakaraensis* TK1769; HBT TRMB, *Hbt. salinarum* VNG 1451C. *Har. hispanica* TrmB homologs are named according to their identity to characterized proteins.

However, the majority of characterized Halobacteria, hereon referred to as haloarchaea for clarity, are carbohydrate utilizers, or saccharolytic. These organisms primarily use the semi-phosphorylative Entner-Doudoroff (spED) pathway for glycolysis, while a modified Embden Meyerhof Parnas (EMP) pathway is predominately utilized during gluconeogenesis and fructose degradation [31–33]. Glycolysis in saccharolytic haloarchaea has two distinguishing features from other archaeal groups that enable the same ATP yield as the classical ED pathway in Bacteria: (i) a bacterial-type catabolic glyceraldehyde-3-phosphate dehydrogenase (GAPDH), and (ii) an amphibolic archaeal phosphoglycerate kinase [34]. Nonsaccharolytic species, such as *Hbt. salinarum* and *Natrinema sp.* strain J7-2, possess all genes necessary for a functional spED pathway except that they lack the *gapI/pgk* operon present in saccharolytic genomes and instead encode an archaeal-specific nonphosphorylating GAP dehydrogenase (GAPN) [35–37]. Despite the presence of all necessary genes in the genome, the activity of spED enzymes has not been detected in *Hbt. salinarum* [24, 27, 38]. In particular, because the GAPN reaction proceeds without the formation of 1,3-bisphosphoglycerate coupled to substrate-level phosphorylation, the theoretical ATP yield of glycolysis in nonsaccharolytic haloarchaea is lower than in haloarchaea possessing a bacterial-type GAPDH.

The role of TrmB has yet to be investigated in other haloarchaea, particularly in models that catabolize diverse sets of carbohydrates and are of interest for industrial applications [39, 40], such as *Haloarcula hispanica* ATCC33960. *Har. hispanica* is a moderately halophilic archaeon that grows on a wide array of carbon sources such as pentose and hexose sugars, disaccharides, three-carbon molecules, and acetate [41, 42]. Moreover, when oxygen, nitrogen, or phosphorus is limited and carbon is abundant, *Har. hispanica* accumulates large quantities of PHBV, a biodegradable plastic alternative [43–45]. Due to its biotechnological potential, metabolic studies thus far have focused on PHBV synthesis [13, 46, 47]. Correct activation of PHBV production implies sensing carbon availability, down-regulation of glycolysis, and activation of the PHBV synthesis pathway. Our understanding of how various signals are integrated to coordinate such a response is still unclear. A better understanding of metabolic regulation generally in *Har. hispanica* is needed to elucidate the function of genes and enzymes for biotechnological applications.

In this study, we characterized TrmB in *Haloarcula hispanica* using high-throughput phenotyping, genome-wide binding assays, and expression experiments to compare its targets with those previously reported in *Hbt. salinarum* [10]. We find that TrmB is essential for growth in gluconeogenic conditions in both species, but when TrmB is deleted in *Har. hispanica* growth can be restored by supplementing a wider variety of carbon sources, in line with its saccharolytic capabilities. We identified 15 robust TrmB binding sites across the genome corresponding to the differential expression of 9 genes predominately involved in gluconeogenesis. A point of bidirectional regulation by TrmB of the EMP pathway in *Hbt. salinarum* (activation of *ppsA* and repression of *pykA*) is absent in *Har. hispanica*. Instead, we propose that TrmB-dependent induction of an archaeal-type GAPDH is necessary for gluconeogenesis. Together these results suggest an ancestral role for TrmB in enabling gluconeogenesis across Euryarchaea and highlight this family of transcriptional regulators as an important indicator of metabolic versatility in hypersaline-adapted archaea.

## Materials and methods

### Media & growth conditions

All *Har. hispanica* strains were derived from *Haloarcula hispanica* ATCC33960 type strain. *Har. hispanica* was routinely grown on rich medium for *Haloferax volcanii* modified to contain 23% basal salts (YPC23) supplemented with 0.1% glucose (w/v) [48]. During glucose limitation experiments, strains were grown in casamino acid media modified to 23% basal salt concentration (Hh-CA). Briefly, 23% basal salts contain, per liter, 184 g of NaCl, 34.5 g of MgSO_4_ *·* 7 H_2_O, 23 g of MgCl *·* 6 H_2_O, 5.4 g of KCl, and 15.3 mM Tris HCl (pH 7.5). If glucose was supplemented, it was added to a final concentration of 0. 1% (w / v) unless otherwise noted. All media were supplemented with uracil (50 *µ*g/ml) to complement the biosynthetic auxotrophy of the Δ*pyrF* parent strain unless specified. Other ingredients and media preparation are in line with Allers *et. al.* [48]. All plates were incubated at 37*^◦^*C for 8-10 days for single colonies and liquid cultures were cultivated aerobically at 37*^◦^*C with 250 rpm orbital agitation.

### Strains, plasmids, & primers

Deletion and integration plasmids were constructed using isothermal assembly [49]. For growth complementation assays, a heterologous expression vector for *Har. hispanica* was constructed from pWL502, a pyrF-based expression vector for *Haloferax mediterranei* [50]. Briefly, a mevinolin resistance cassette was amplified from pNBKO7 and replaced the *Hfx. mediterranei pyrF* at the SmaI and BamHI sites to yield pAKS83. All plasmid sequences were confirmed via Sanger sequencing and propagated in *Escherichia coli* NEB5*α*. Strains, plasmids, and primers used are presented in S1 Table, S2 Table, S3 Table, respectively. Gene deletions and chromosomal integrations were performed using two-stage selection and counterselection as described previously [44]. Strains were generated using the spheroplasting transformation method and plated on Hh-CA without supplemental uracil [51]. Resulting colonies were inoculated in 5 ml of YPC23 + glucose and grown for 48 hours and then plated onto Hh-CA + 150 *µ*g/ml 5-Fluoroorotic acid (5-FOA) + glucose for counterselection.

To verify the complete deletion of all copies of *trmB* in the genome and to check for secondary mutations, genomic DNA was extracted using phenol:chloroform:isoamyl alcohol (25:24:1) followed by ethanol precipitation. TruSeq libraries were prepared and sequenced on Illumina MiSeq by the Center for Genomic and Computational Biology at Duke University (Durham, NC). Reads were aligned to the *Har. hispanica* ATCC33960 genome (GCF 000223905.1, assembly ID ASM22390v1, accessed 2022-10-19) using breseq with default options [52].

### Sequence analysis of *Har. hispanica* TrmB homologs

To identify sugar-sensing TrmB homologs, reference proteomes were searched for protein sequences similar to TrmB_Hbt_ and results were filtered to include hits with identical domain architecture. Domain identities and confidence values were confirmed using hmmscan on the Pfam database [53]. *Har. hispanica*TrmB protein and gene sequences were accessed from NCBI and globally aligned to *Halobacterium salinarum NRC-1* VNG1451C using EMBOSS Needle [54].

### Quantitative phenotyping

Strains were struck from freezer stocks for each experiment and incubated for 10 days. Single colonies were inoculated in 5 mL YPC23 + glucose and grown to stationary phase (OD_600_ *∼* 4). Stationary precultures were then collected, washed twice in Hh-CA and diluted into 200 *µ*l fresh Hh-CA with or without 25 mM of additional carbon sources to an initial OD_600_ of 0.05. Growth was measured every 30 minutes at 37*^◦^*C with continuous shaking for 72-90 hours in Bioscreen C analysis system (Growth Curves USA, Piscataway, NJ). For quantitative analysis, growth curves were blank-adjusted within independent experiments, fitted, and holistic differences across growth phases were summarized using the area under the curve (AUC) [55]. AUC values were averaged across technical replicates, the standard deviation was calculated across biological replicates, and significant differences were evaluated by a two-tailed paired Student’s t-test.

### Isolation of uracil prototrophic AKS133 strains

Colonies used for AKS133 RNA-seq experiment that showed *pyrF* expression were subjected to a quantitative growth assay in Hh-CA media *±* uracil *±* glucose to test for growth in the absence of uracil. All 10 wells containing AKS133 grew in the absence of uracil. These cultures were inoculated into Hh-CA - uracil + glucose for 24 hours and then plated in medium lacking uracil for individual colonies. Of the ten cultures, single colonies were isolated from 7, from which genomic DNA was extracted. Amplification of *pyrF* from genomic DNA confirmed that the endogenous deletion was intact in all prototrophic isolates, ruling out gene conversion, and revealed *pyrF* sequence elsewhere in the genome (primer sequences are listed in S3 Table). We were unable to locate *pyrF* with multiple attempts at arbitrary PCR [56], but note that there is substantial sequence homology between the vector and the *Har. hispanica* genome, particularly near the origins of replication.

### AKS319 strain construction & verification

To construct AKS319, transformations were carried out as described above, except that all media and plates were supplemented with glucose to alleviate selection. Colonies harboring the deletion vector were passaged twice in YPC23 + glucose (for a total of 96 hours), and then exposed to a higher concentration (250 *µ*g/ml) of 5-FOA to select against *pryF* and vector retention. After initial confirmation of genotype by PCR, prior to storage and WGS, 5-FOA-resistant colonies were screened for uracil prototrophic growth for 90 hours in Hh-CA media *±* uracil *±* glucose. Genomic DNA was extracted and WGS was carried out to confirm no reads mapped to the *pyrF* or *trmB* loci.

### Chromatin immunoprecipitation

Strains DF60 and AKS155 were struck from freezer stocks onto YPC23 plates and incubated for 8 days. Independent colonies were used to start two cultures of DF60 and four cultures of AKS155 in 10 mL YPC23. Precultures were grown for 29 hours (OD_600_ *∼* 2) then collected at 6000 rpm for 2 minutes and washed twice with Hh-CA. AKS155 pellets were resuspended in Hh-CA or Hh-CA + glucose and grown to midlog phase (average OD_600_ *∼* 0.34) before crosslinking (Supplementary File 1). Samples were processed as described by Wilbanks et al. using anti-HA polyclonal antibody (Abcam catalog #ab9110) to immunoprecipitate cross-linked fragments [57]. Libraries were constructed by the Center for Genomic and Computational Biology at Duke University (Durham, NC) using KAPA Hyper Prep kit and Illumina TruSeq adapters, and the 50 base pair, single-ended libraries were sequenced on an Illumina HiSeq 4000.

### ChIP-seq read processing & peak calling

Raw FastQ files were trimmed of adapter sequences with Trim galore! 0.4.3 and Cutadapt 2.3 and read quality was checked with FastQC 0.11.7. Reads were aligned to the *Har. hispanica* genome with Bowtie2 2.3.4.3 [58]. Aligned sequence files were then sorted, indexed, and converted to binary format with samtools 1.9 [59]. Before calling the peaks, the fragment length was optimized for each IP and input control sample using ChIPQC 1.30 [60]. Binding peaks were called from sorted bam files using Mosaics 2.32 in R 4.1.2 with the calculated fragment lengths and FDR *<* 0.01. DiffBind 3.4.11 was used to merge peaks that were shared in at least 3 biological replicates for samples grown without glucose (N=4) and all samples grown in the presence of glucose (N=2) [61]. Briefly, binding peaks were merged if they overlapped by at least 1 base pair and consensus peaks were then trimmed to 300 base pair width centered around the consensus peak maximum. Samples were RLE normalized. Separately, we used DiffBind to identify peaks shared between glucose-replete and depleted samples that exhibited differential binding. Peaks were visually verified using the trackvieweR package to determine an enrichment cutoff [62]. Then, consensus peaks were annotated using IRanges and GenomicFeatures to identify genomic features overlapping and adjacent to TrmB binding peaks [63]. For the per base enrichment plot (Fig 3B), BAM files were extended to their fragment size (150 base pairs), converted to BEDGRAPH format, and scaled according to sequencing depth before calculating the enrichment ratio and averaging across replicates as described by Grünberger *et al.* [64].

### RNA-isolation & sequencing

Strains DF60 and ASK133 or AKS319 were struck from freezer stocks onto YPC23 + 0.1% glucose plates and incubated for 10 days. Four single colonies of DF60 and ASK133 or AKS319 were inoculated in 5 mL YPC23 medium with 0.1% glucose and grown aerobically to stationary phase (OD_600_ *∼* 4). Cells were washed twice with Hh-CA medium without glucose and transferred into fresh 50 mL Hh-CA with or without 0.1% glucose at an initial density of OD_600_ *∼* 0.4. After 24 hours, cultures were harvested and RNA extracted using Absolutely RNA Miniprep kit (Agilent Technologies, Santa Clara, CA) followed by additional DNAse treatment with Turbo DNAse (Invitrogen, Waltham, MA). Total RNA was quantified using Agilent Bioanalyzer RNA Nano 6000 chip (Agilent Technologies, Santa Clara, CA). The absence of DNA contamination was determined on 200 ng of RNA in 25 PCR cycles. Ribosomal RNA was removed with the PanArchaea riboPOOL kit according to the manufacturer protocol (siTOOLs Biotech, Germany), and sequencing libraries were constructed with NEBNext UltraII Directional RNA Library Preparation Kit (Illumina, #E7760) as described previously [65]. The fragment size of the libraries was measured using the Agilent Bioanalyzer DNA 1000 chip and then pooled and sequenced on NovaSeq6000 at the Center for Genomic and Computational Biology at Duke University (Durham, NC).

### Differential expression analysis & clustering

FastQ files were processed as described for ChIP-seq, except alignment by Bowtie2 required stand-specific read orientation. After alignment, counts were calculated using featureCounts requiring complete and concordant alignment by paired reads [66].

Samples were considered outliers and removed prior to differential expression analysis if they had a pairwise correlation lower than R=0.6 with all other replicates. Differential expression analysis was carried out in DeSeq2 with logarithmic change and FDR cutoffs of 1 and 0.05, respectively, using an experimental design informed by TrmB model of regulation in *Hbt. salinarum*[10, 67]. To cluster differentially expressed genes based on expression patterns across strain and condition, normalized counts were mean and variance standardized and subjected to k-means clustering and visualized as described in [68].

### Functional enrichment

Annotated and predicted gene functions were accessed from EggNOG 5.1 [69]. Genes adjacent to TrmB binding sites were then tested for functional enrichment relative to the whole genome using a hypergeometric test with Benjamini and Hochberg multiple testing correction.

### Motif discovery & scanning

TrmB binding motif was identified using the command line version of MEME Suite 5.5.1. Specifically, sequences under consensus peaks (300 base pairs) were extracted and analyzed with XSTREME for motif discovery, motif enrichment, and motif search [70]. Default settings were used except that the maximum motif width was set to 19 base pairs and a first-order Markov background model was generated using non-coding regions of the *Har. hispanica* genome to control for dinucleotide frequencies. The recognition motif reported for *Hbt. salinarum* was included for downstream motif enrichment and similarity assessment (Schmid *et. al.* supplemental table 5). The motif identified was robust to zero and second-order background models. To identify motif occurrences genome-wide, the *Haloarcula hispanica* reference genome was scanned using FIMO with a p-value cutoff of 1 *×* 10*^−^*^4^.

### Data availability

All code used to analyze data and generate the figures presented here can be found on Git Hub (https://github.com/hackkr/Har_hispanica_TrmB). Whole genome sequencing data is available in the National Center for Biotechnology Information (NCBI) Sequence Read Archive under PRJNA947196. RNA-seq and ChIP-seq raw data, counts, and coverage files are available on Gene Expression Omnibus under accession number GSE227034.

## Results

### Four TrmB homologs are encoded in the *Har. Hispanica* genome

Using bioinformatic analysis, we investigated the conservation of TrmB. We detected 619 proteins with identical domain architecture as *Hbt. salinarum* TrmB across archaeal and bacterial Pfam reference proteomes [71]. Of these, almost half were encoded in haloarchaeal genomes (Fig 1A). On average, Haloarchaea genomes contain three TrmB homologs per genome, compared to an average of one homolog outside of haloarchaea, indicating lineage-specific expansion of this class of regulators. Some regulators containing the TrmB DNA-binding domain have been recently shown to function during oxidative stress in haloarchaea [22], but proteins with both a TrmB DNA-binding domain and carbohydrate-binding domain have not been functionally characterized in haloarchaea aside from VNG 1451C in *Hbt. salinarum* (hereafter referred to as TrmB_Hbt_).

To determine the most likely functional ortholog of *Hbt. salinarum* TrmB (VNG 1451C) in the *Har. hispanica* genome, we searched the 148 proteins with predicted DNA-binding capabilities. Ten of these putative transcription factors have a TrmB DNA-binding domain and four contain both the TrmB DNA-binding domain and sugar-binding domain (PF01978 and PF11495, respectively): HAH 0923, HAH 1548, HAH 1557, and HAH 2795 (Fig 1B). Multiple sequence alignment revealed that HAH 0923, HAH 1548, and HAH 2795 maintain two of the six active site residues required for sugar-binding in *Thermococcus litoralis*, while HAH 1557 preserved four of the six residues [72]. Both G320 and E326 residues, which when mutated in *Tcc. litoralis* drastically reduce the binding affinity for sucrose and maltose, are conserved across the haloarchaeal TrmB proteins analyzed [72]. In contrast, N305, which is specific for maltose but not sucrose binding, is not conserved outside of hyperthermophiles [73]. Of the TrmB homologs in *Har. hispanica*, HAH 1548 exhibits the highest amino acid sequence identity to TrmB_Hbt_, and best agreement with the sugar-binding domain consensus (e-value = 3.3 *×* 10*^−^*^20^, Table 1).

**Table 1.** *Har. hispanica* TrmB homologs.

| TrmB homolog | gene name | protein identity | protein positives | DBD e-value | SBD e-value |
| --- | --- | --- | --- | --- | --- |
| VNG_1451 | *trmB* | 100% | 100% | $1.8 \times 10^{-70}$ | $6.1 \times 10^{-20}$ |
| **HAH_1548** | *trmB* | **56.5%** | **74.2%** | $1.1 \times 10^{-68}$ | $3.3 \times 10^{-20}$ |
| HAH_2795 | *trmB2* | 31.2% | 47.5% | $1.5 \times 10^{-70}$ | $3.5 \times 10^{-14}$ |
| HAH_1557 | *trmB3* | 27.2% | 48.5% | $2.9 \times 10^{-78}$ | $1.2 \times 10^{-14}$ |
| HAH_0923 | *trmB4* | 25.4% | 39.9% | $1.9 \times 10^{-60}$ | $4.0 \times 10^{-17}$ |

Protein sequences were globally aligned to *Hbt. salinarum* TrmB. DNA-binding domain (DBD) and sugar-binding domain (SBD) confidence scores were calculated with hmmscan [53].

In *Tcc. litoralis* and *Tcc. kodakarensis*, the *trmB* homolog is located near genes that encode carbohydrate ABC transporters. However, in *Hbt. salinarum* and *Pyr. furiosus*, these genes are absent from the genomic region surrounding *trmB* [10, 74]. Genomic context analysis revealed that HAH 1548 is upstream of a putative carbohydrate ABC transporter and glucose-1-dehydrogenase. HAH 1557 flanks the same ABC transport system but has a substantially lower identity to TrmB_Hbt_ (Fig. 1B, Table 1). Thus, HAH 1548 was distinguished via sequence alignment and genomic context as the likely functional homolog of TrmB_Hbt_ (for clarity, hereafter we refer to the HAH 1548 gene as *trmB_Har_* and the translated protein as TrmB_Har_).

### TrmB_Har_ is essential for growth in gluconeogenic conditions

To assess whether TrmB_Har_ plays a physiological role in carbohydrate metabolism and gluconeogenesis, we deleted *trmB_Har_*. Because haloarchaea are polyploid [75], it is possible for copies of the wild-type locus to persist at levels too low to detect via Sanger sequencing [76]. Therefore, the genotypes of all strains in this study were confirmed with short-read whole genome sequencing (WGS). WGS of the *trmB_Har_* deletion strain confirmed it to be free of second-site mutations and chromosomal copies of *trmB_Har_* (S1 Fig, SRA accession PRJNA947196). Notably, DF60, the Δ*pyrF* parent strain, harbors a previously unreported 2.6 kb deletion disrupting HAH 2675-79, encoding *flgA1* and *cheW1*. This region is absent in all strains derived from the Δ*pyrF* strain in this study.

We subjected both Δ*trmB_Hbt_* and Δ*trmB_Har_* to a panel of nine ecologically and physiologically relevant carbon sources at an equimolar concentration to facilitate comparisons between species (Fig. 2). As expected from previous reports, the *Hbt. salinarum* Δ*trmB* strain exhibits a severe growth defect under gluconeogenic conditions [10]. Only glucose and glycerol stimulate growth in the deletion strain (Fig 2A). The slight differences between these results and those reported by Schmid *et al.* are due to differences in supplemental carbon concentration (389mM and 760mM for glucose and glycerol, respectively, [10]).

**Fig 2.**
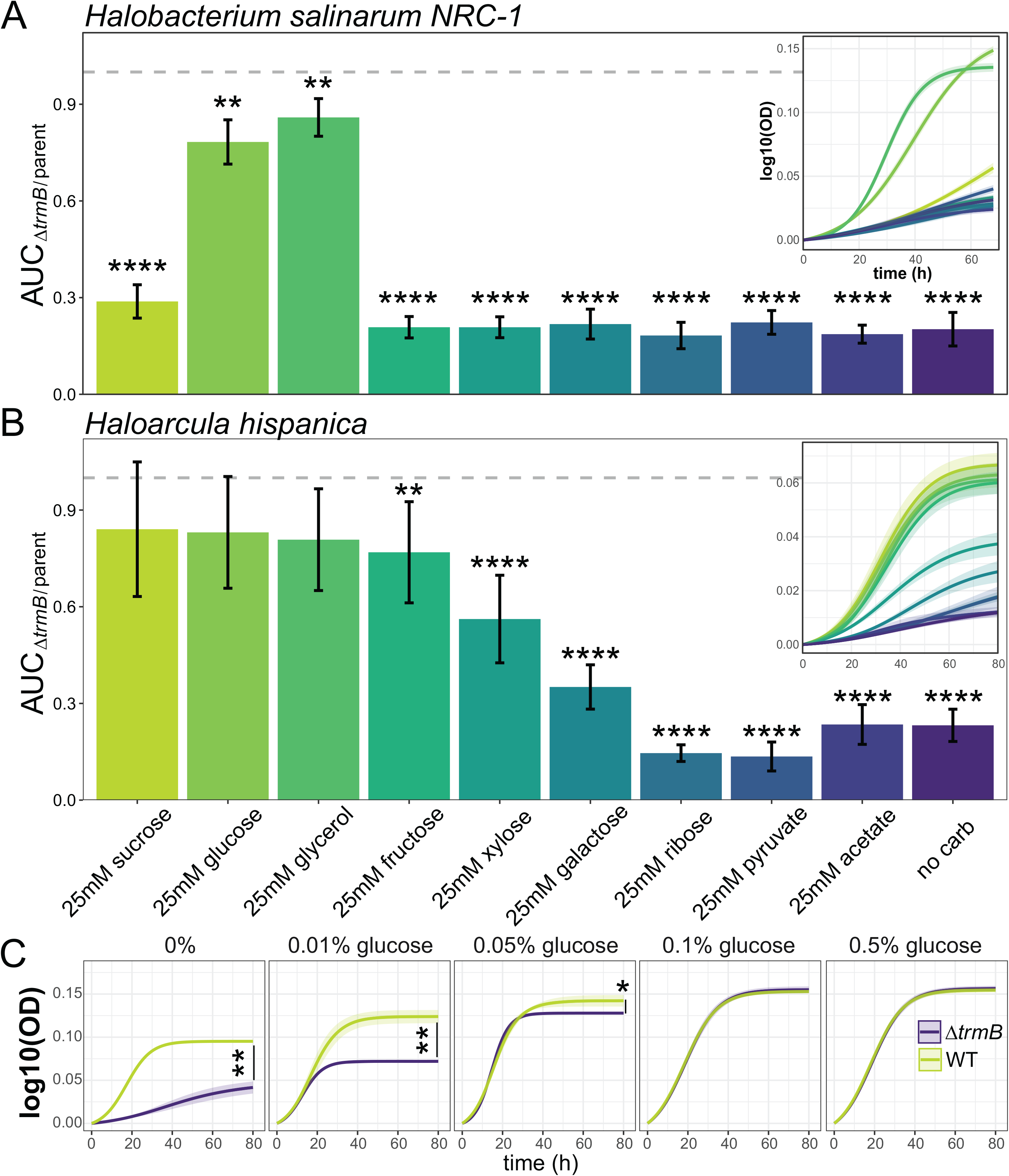
TrmB_Har_ is essential for gluconeogenic growth. Proportion of parent growth achieved by (A) Δ*trmB_Hbt_* and (B) Δ*trmB_Har_* with various carbon supplements, as measured by area under the growth curve (AUC). Error bars indicate the standard deviation over a minimum of 3 biological replicates, each in technical triplicate. Insets show the fitted, log-transformed growth curves of Δ*trmB* strains in each condition. C) Effect of increasing glucose concentrations on the growth of *Har. hispanica* Δ*trmB* and Δ*pyrF* strains. Solid lines represent the mean of the log-transformed growth curve, and shaded regions depict 95% confidence intervals. For all panels, asterisks indicate significance: * *p*-value *<* 0.05; ** *<* 0.01; **** *<* 0.0001.

Growth assays also revealed a severe growth defect of the Δ*trmB_Har_* strain in Hh-CA (Fig 2B). Without additional carbon sources, amino acids are the sole nitrogen and carbon source in Hh-CA, so cells rely on gluconeogenesis to synthesize all required glucose. Normal growth could be restored with heterologous expression of *TrmB_Har_* (S2 Fig). In contrast to *Hbt. salinarum*, Δ*trmB_Har_* growth could be restored by the addition of glucose, fructose, sucrose, or glycerol. Xylose stimulated growth to 57% of the parent strain, as measured by the area under the growth curve (AUC), but growth remains depressed relative to the parent strain in the presence of ribose, pyruvate, acetate, and galactose. Accelerated growth of the parent strain in media supplemented with pyruvate and ribose contributes to the lower growth ratios reported for those carbon sources (S2 Fig).

We next titrated the concentration of glucose to identify the minimum amount that restored normal growth (Fig 2C). Relative to parent strain growth in the absence of carbon, Δ*trmB_Har_* was stimulated 0.31, 0.81, 1.42, 1.73, 1.74-fold with no supplemental carbon source, 0.01%, 0.05%, 0.1%, and 0.5% glucose (w/v), respectively. Glucose concentrations above 0.1% did not further stimulate growth, identifying 0.1% glucose, or 5.55 mM, as the minimal glucose concentration needed to rescue growth. These data demonstrate that TrmB is indispensable for growth under gluconeogenic conditions in haloarchaea with various metabolic capabilities, and implicate glucose in the function of TrmB in both *Hbt. salinarum* and *Har. hispanica*.

### TrmB_Har_ binds promoters of genes involved in central carbon metabolism

To determine whether TrmB_Har_ functions as a transcriptional regulator, we located TrmB_Har_-DNA interactions with chromatin immunoprecipitation coupled to sequencing (ChIP-Seq) in the presence and absence of glucose. A *trmB_Har_*::hemagglutinin (HA) epitope fusion at the endogenous locus was constructed for these experiments and confirmed via WGS (S1 Fig). There was no difference in the growth of the tagged strain from the parent strain in the absence of glucose, suggesting the C-terminal fusion retained full functionality (S2 Fig).

Consistent with the model of regulation reported in *Hbt. salinarum* and homologs in hyperthermophiles [10, 17, 77], TrmB_Har_ primarily binds DNA when glucose is absent (Fig 3A). Under this condition, fifteen reproducible TrmB_Har_ binding sites, or peaks, were enriched relative to the input control (Table 2, Fig 3B, Methods). Of the 15 peaks, seven exhibited significantly increased binding in the absence of glucose relative to glucose-replete samples (FDR *≤* 0.1). Seven other peaks were exclusively detected in the absence of glucose, and one was observed in all samples regardless of glucose availability (S1 File, S3 Fig).

**Table 2.**
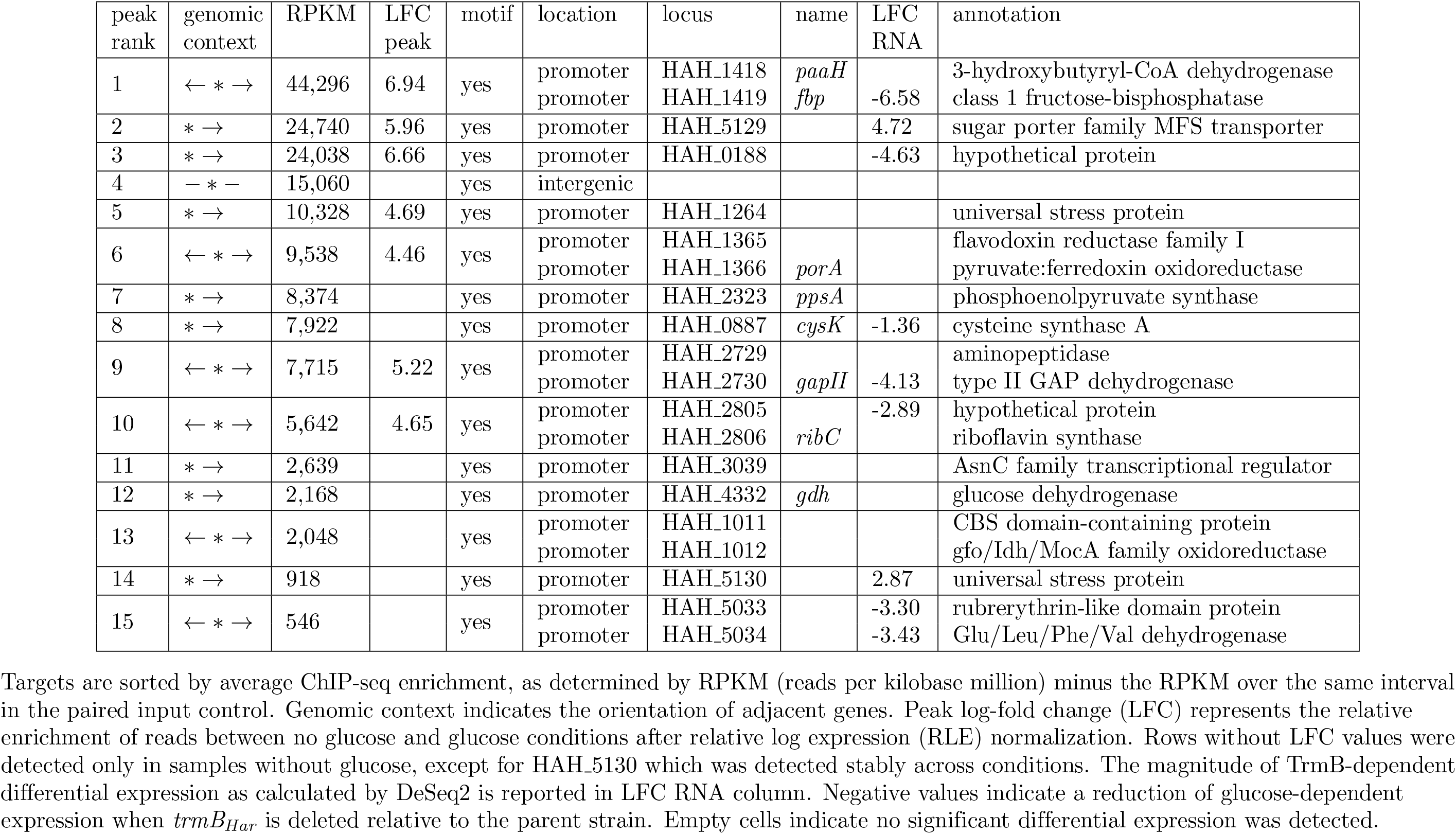
*Har. hispanica* TrmB binding-sites.

**Fig 3.**
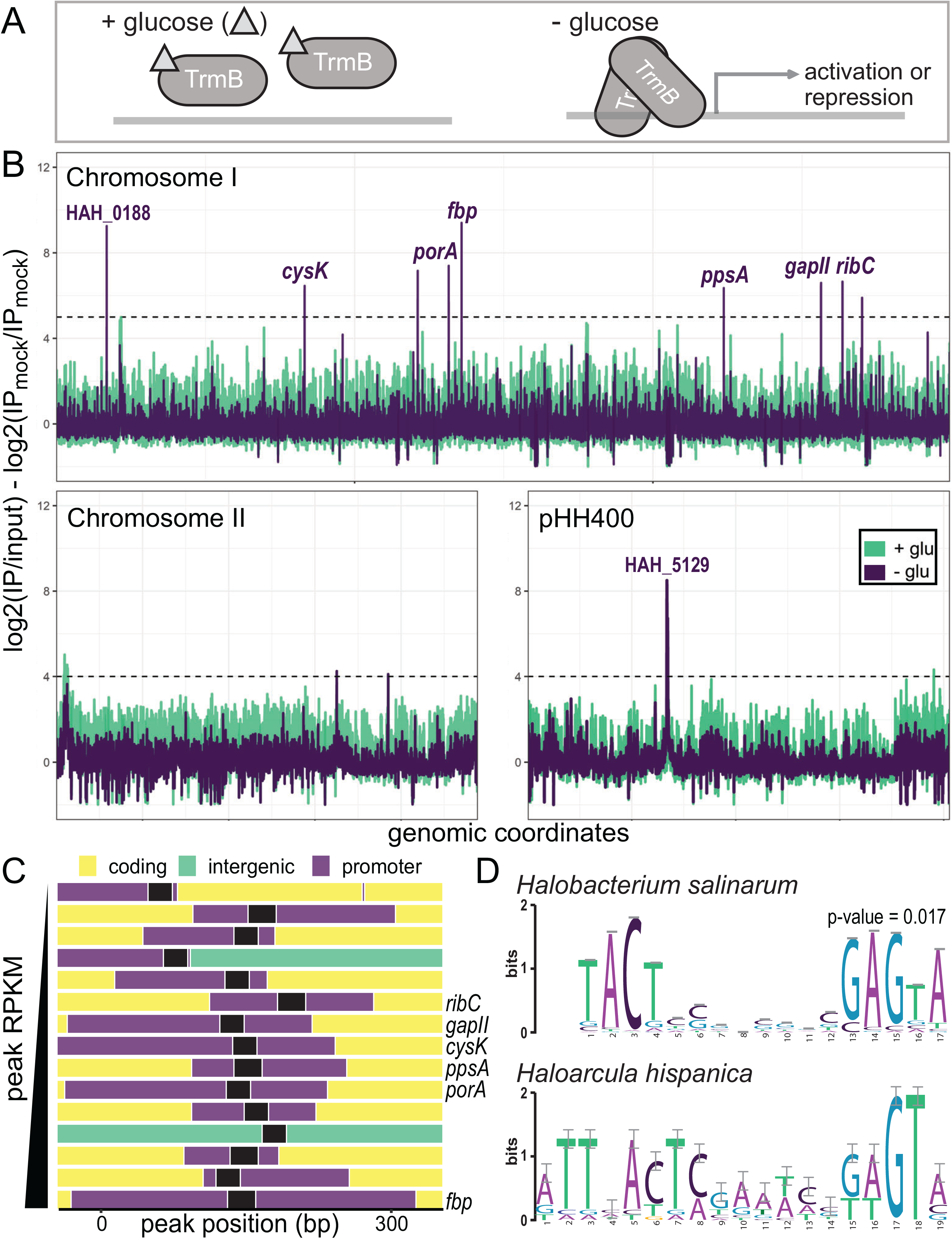
Genome-wide TrmB_Har_ binding sites. A) Proposed TrmB mechanism of regulation from *Hbt. salinarum* and *Thermococcus kodakaraensis* [10, 19]. B) Per base enrichment of reads across the genome. Samples were normalized to paired input samples and averaged across replicates. Averages were normalized relative to untagged controls and log-transformed. C) Genomic context of TrmB_Har_ binding sites. Sites were trimmed to 300 base pairs centered around peak summits. Promoter regions are shown in purple, intergenic regions in green, coding regions in yellow, and motif locations in black. Regions are sorted by enrichment, or peak size. D) XSTREME motif analysis of selected regions. The 19-base pair motif identified is similar to the binding motif reported for TrmB_Hbt_ (SEA p-value = 0.017).

TrmB_Har_ binding sites were predominantly located in non-coding regions of the genome (Fig 3C): 41% of the bases in enriched regions were non-coding (p-value *<* 1 *×* 10*^−^*^15^, binomial test), despite non-coding sequences comprising only 13% of the *Har. hispanica* genome. Because archaeal core promoter consensus sequences are still ill-defined, the non-coding regions up to 250 base pairs upstream of translational start sites were considered promoter regions. TrmB_Har_ binding sites overlap the promoter regions of 20 genes (Table 2). Of those, five are homologous to TrmB_Hbt_ targets by reciprocal protein blast, all in the canonical EMP gluconeogenic pathway: pyruvate oxidoreductase (encoded by *porA/B*), phosphoenolpyruvate synthase (*ppsA*), GAPDHII (*gapII*), and fructose-bisphosphatase (*fbp*). Overall, the predicted function of genes near binding sites were significantly enriched for carbohydrate transport and metabolism (adjusted p-value *<* 7.3 *×* 10*^−^*^5^, hypergeometric test). Notably, the second largest peak is located in the promoter of HAH 5129, a sugar major facilitator superfamily (MFS) transporter (PF00083, e-value *<* 3.0 *×* 10*^−^*^123^), highlighting this gene as a candidate for the primary glucose transporter in *Har. hispanica*.

Computational *de novo* motif detection using 300 base pairs around binding summits revealed a 19 base pair palindromic binding motif (Fig 3D). The motif is similar to the TrmB recognition sequence reported for *Hbt. salinarum* (p-value = 0.017, calculated with TomTom [78]), and robust to various background models (S4 Fig). The motif occurs 235 times throughout the genome, and of those, 67 instances occur in the promoter regions of 58 genes (motif occurrences on opposite strands were considered distinct, S1 File). Like genes near peaks, genes with motifs located in promoter regions are enriched for functions in the transport and metabolism of carbohydrates and amino acids (hypergeometric test adjusted p-value *<* 5.2 *×* 10*^−^*^5^ and 0.003, respectively).

Together, these data support the model that in the absence of glucose TrmB_Har_ recognizes a conserved cis-regulatory sequence to bind DNA near genes primarily involved in carbon metabolism.

### Transcriptome profiling suggests cryptic integration of deletion vector, resulting in uracil prototrophy in **Δ***trmB_Har_* strains

We conducted transcriptome profiling to determine the direction of TrmB-dependent regulation of genes near binding sites and to elucidate TrmB_Har_ targets in instances where peaks were located over the promoters of divergent genes. Unexpectedly, we observed high *pyrF* expression even though the *pyrF* gene (the *ura5* homolog) is deleted in the parent strain to enable genetic manipulation [44]. Transcripts mapping to the *pryF* locus were not present in any parent samples but were detected in all Δ*trmB_Har_* (AKS133) samples regardless of glucose availability. Subsequent phenotyping and genotyping confirmed stably inherited uracil prototrophy in 7 of 10 independent AKS133 isolates and ruled out stock contamination, vector propagation, and gene conversion at the endogenous locus (S5 Fig; Methods). This points to homologous recombination of the deletion vector, which harbors *pryF*, elsewhere in the genome.

We carefully generated a new Δ*trmB_Har_* strain, AKS319, and repeated the experiment. Despite additional safeguards during strain generation and genotype confirmation by WGS, 4 of 8 Δ*trmB_Har_* samples exhibited high *pyrF* expression (S6 Fig; Methods). Due to this variation, however, we were able to investigate the fitness and transcriptomic consequences of *pyrF* expression in the AKS319 strain and across Δ*trmB_Har_* strains (S6 Fig). AKS319 exhibits a more extreme growth defect relative to AKS133, which can be mostly, but not completely, ameliorated by glucose supplementation. However, expression of *pyrF* has little effect on the overall transcriptome: average counts per gene are highly correlated across all Δ*trmB_Har_*samples regardless of *pyrF* expression both before (S6 Fig) and after batch correction(Fig 4A, S1 File). The repeated and apparently independent instances of vector integration resulting in uracil prototrophy suggests that both deletion strains are likely mixed populations with a small proportion of cells retaining vector sequence.

**Fig 4.**
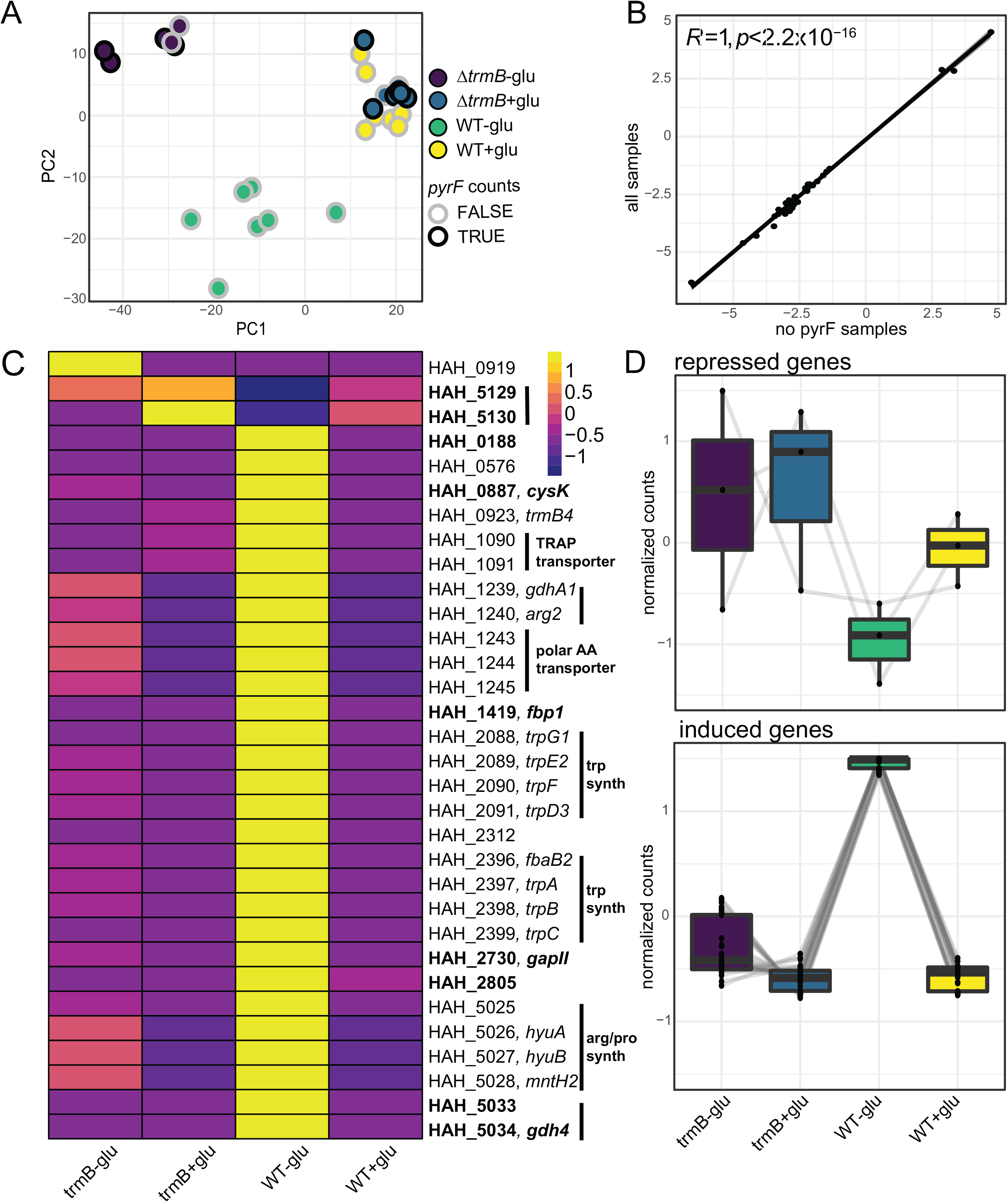
TrmB_Har_-dependent expression. A) Principal component analysis of AKS133 and AKS319 samples after batch correction. Samples with transcripts mapping to *pyrF* are outlined in black. B) Correlation of 32 differentially expressed genes from analyses including all Δ*trmB_Har_* samples (y-axis) and only samples with no detected *pyrF* expression (x-axis). C) Normalized expression pattern of 32 genes in (B). Yellow indicates elevated expression and purple indicates reduced expression. Genes near peaks are bolded and genes predicted to be co-transcribed are indicated by a black bar. If available, the predicted function of the operon was provided. D) Three and 29 genes have patterns consistent with TrmB-dependent repression and induction, respectively. Light grey lines connect normalized expression values for each transcript across strain and condition.

TrmB_Har_ may be essential in the DF60 background, explaining this phenomenon.

### TrmB_Har_ activates genes involved in gluconeogenesis and tryptophan biosynthesis and represses those encoding glucose uptake

Based on the model of TrmB regulation described in *Hbt. salinarum*, we reasoned that direct targets of TrmB_Har_ would exhibit differential expression only when TrmB is present and active, i.e., in the parent strain grown in the absence of glucose (Fig 3A). This assumption was made explicit in the DeSeq2 model used to identify differentially expressed genes [67]. Using this framework, we detected 32 genes that were significantly differentially expressed (FDR *<* 0.05; LFC *>* 1) in the parent vs mutant strain across two independent analyses, one including and the other excluding samples with counts mapping to *pyrF* (Fig 4B, S1 File). Because *pyrF* expression, when present, is not glucose-dependent, it was not considered significant using this explicit model.

Of the 32 genes, three are significantly up-regulated in Δ*trmB_Har_* relative to the parent strain in the absence of glucose: glutamate synthase (encoded by *HAH 0919*), a universal stress protein (*HAH 5129*), and the MFS family sugar transporter (*HAH 5130*) (Fig 4C and D). The remaining 29 genes exhibit the opposite pattern: they are significantly down-regulated in the deletion strain, exhibiting a pattern across strains and conditions consistent with TrmB-mediated induction. The predicted products of these 29 genes are enriched for functions in amino acid transport and metabolism (adjusted p-value = 1.085 *×* 10*^−^*^10^, hypergeometric test) including two tryptophan biosynthesis operons, cysteine synthase, a putative amino acid transporter, and a TRAP transporter (Fig 4D). Genes whose expression is induced also include those predicted to function in gluconeogenesis (encoding an archaeal class II GAPDH, class I fructose-bisphosphate) and two small hypothetical proteins less than 50 amino acids in length (*HAH 0188* and *HAH 2805*).

Integrating across experiments, nine genes are adjacent to TrmB_Har_ binding sites containing a palindromic cis-regulatory motif, and are differentially expressed (Table 2, Fig 3D). Taking the ChIP-seq, RNA-seq, and motif evidence together, we conclude that these genes comprise the high confidence regulon under the direct transcriptional control of TrmB_Har_ (Fig 4C bolded, Fig 5A). For these targets, we also determined whether the relative TrmB motif distance from the putative TATA box or start codon was predictive of the direction of regulation (S4 Table). The archaeal promoter architecture resembles a simplified version of that found in eukaryotes, including a TATA box located around -26 bp and a BRE element around -33 bp [79, 80]. Generally, motifs were upstream of predicted transcription initiation features, consistent with our observation of TrmB acting as an activator of these genes. In contrast, genes repressed by TrmB binding had motifs overlapping the predicted B recognition elements.

**Fig 5.**
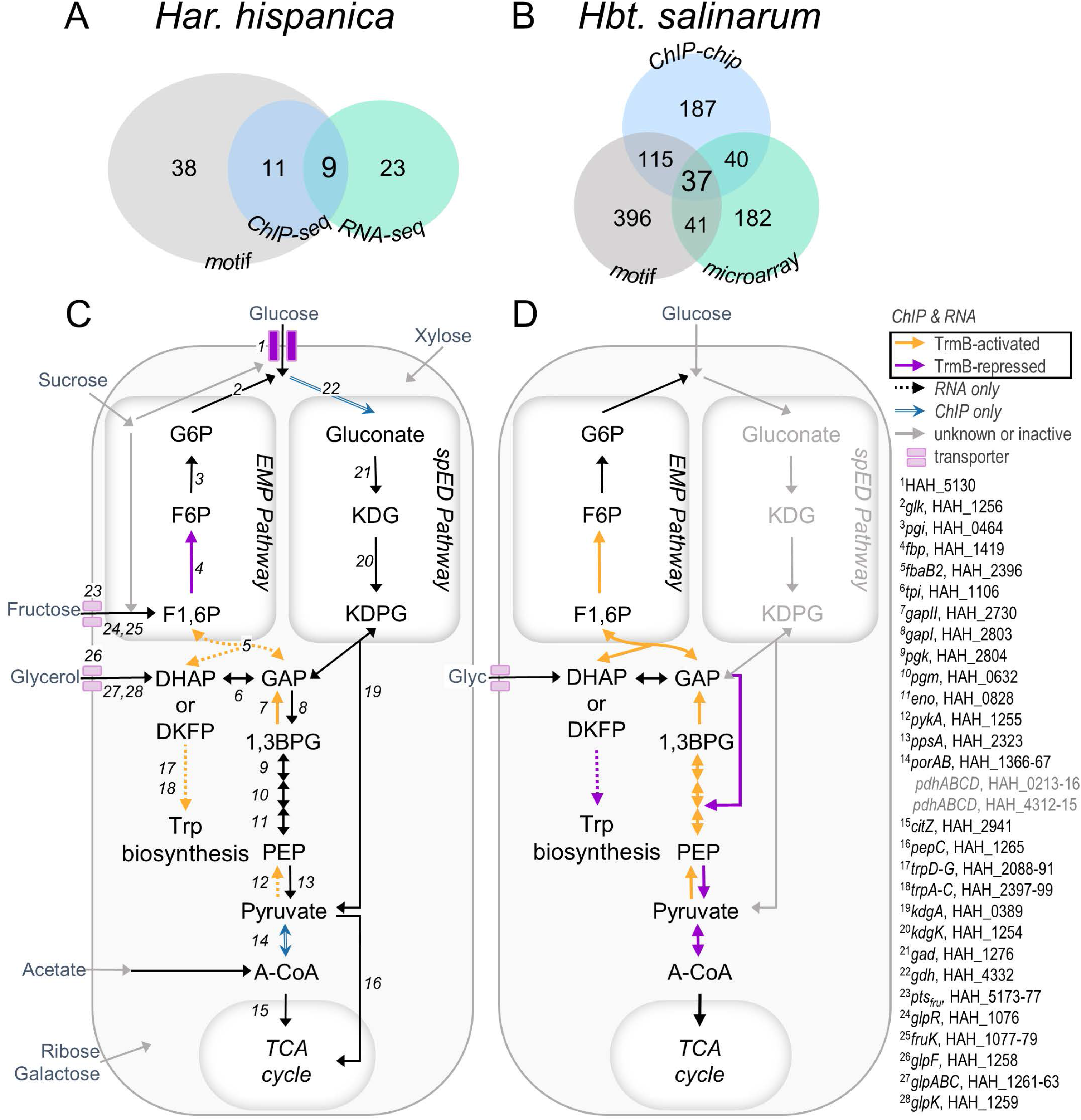
Comparison of TrmB targets in gluconeogenesis in haloarchaea. Summary of experimental evidence of TrmB regulon in (A) *Har. hispanica* and (B) *Hbt. salinarum*. Data for *Hbt. salinarum* summarized from figures 4 and 6 of Schmid *et. al.* [10]. C, D: Summary of TrmB targets that encode enzymes in central carbon metabolism for each species. Targets in the high-confidence regulon are indicated with solid, colored arrows. Genes differentially expressed but lacking a nearby binding site are indicated with dashed arrows. Targets that were near binding sites but not differentially expressed are indicated in light blue. Solid gray arrows indicate that specific carbohydrate transport systems are unknown or that genes are present but no enzymatic activity has been detected. Genes predicted to encode the necessary enzymes are labeled (see S8 FigA for normalized expression values). Additional putative pyruvate oxidation systems that are not regulated by TrmB are listed in gray. G6P, glucose-6-phosphate; F6P, fructose-6-phosphate; F1,6P, fructose-1,6-bisphosphate; DHAP, dihydroxyacetone phosphate; DKFP, 6-deoxy-5-ketofructose-1-phosphate ; GAP, glyceraldehyde-3-phosphate; 1,3BPG, 1,3-bisphosphoglycerate; PEP, phosphoenolpyruvate; A-CoA, acetyl coenzyme A; KDG, 2-keto-3-deoxygluconate; KDPG, 2-keto-3-deoxy-6-phosphogluconate.

Given the small number of TrmB binding sites across the genome, it was surprising that only half of the binding sites were near genes that exhibited differential expression. We wondered if there might be other regulatory mechanisms at play, namely TrmB_Har_-dependent regulation of small or antisense RNAs, which have recently been described in other haloarchaeal species [81–84]. Since we required correct strand orientation when generating transcript counts, reads mapping to the non-coding strand would not have been considered in the downstream differential expression analysis. We visualized strand-specific reads near peaks and identified an unannotated, monocistronic transcript approximately 200 base pairs in length near the sole intergenic peak between convergently transcribed genes in our data. This transcript is induced when TrmB is active and contains a predicted promoter sequence upstream (S7 Fig). We also saw evidence for a transcript anti-sense to HAH 0887 that appears to be induced when TrmB is active, and that both genes encoding putative small proteins (HAH 0188 and HAH 2805) appear to have extended 3’ UTRs (S7 Fig). We did not find any evidence for unannotated transcripts for peaks near HAH 1010, HAH 1264, HAH 3039, or HAH 4332. The unannotated transcripts require validation and further characterization to be considered direct targets of TrmB_Har_, but support the hypothesis that TFs may regulate the expression of regulatory RNAs in haloarchaea, as has been reported for methanogens [85].

## Discussion

Our data facilitate the general comparison of the TrmB regulon in Euryarchaea and more closely between *Hbt. salinarum* and *Har. hispanica*. TrmB_Har_ has fewer binding sites than TrmBL1 in *Pyr. furiosus* across the genome (15 and 28, respectively [17]). Moreover, TrmB_Har_ acts primarily as an activator of gluconeogenic genes while TrmBL1 was predicted to repress most of its targets, which were involved in glycolysis (21, based on relative motif location) [17]. Similarly to TrmBL1, however, our *in vivo* data suggest that expression of TrmB_Har_ is not autoregulated, contrary to reports of other homologs *in vitro* [10, 86]. Conserved regulon members include *gapII* and an MFS transporter, which are activated and repressed, respectively.

At the sequence level, only two homologs are near TrmB binding sites and exhibit congruent patterns of differential expression in both haloarchaeal species: *fbp* and *gapII* (Fig 5C and D). In general, much more of the triose portion of the EMP pathway (sometimes referred to as ”lower glycolysis”, although here ”upper gluconeogenesis” would be more appropriate) appears to be regulated by TrmB in *Hbt. salinarum* relative to *Har. hispanica* (Figure 5). For example, TrmB represses and activates genes that encode pyruvate kinase and phosphoenolpyruvate synthase, respectively, in *Hbt. salinarum*. In *Har. hispanica*, this control point appears to be less important. This may be because multiple pyruvate oxidation systems are present (listed in 5), including pyruvate carboxylase, homologous to the preferred anaplerotic enzyme in another saccharolytic haloarchaeal species, *Haloferax volcanii* [87]. None of the genes encoding pyruvate oxidation systems were differentially expressed in Δ*trmB_Har_* (S8 FigA), though TrmB_Har_ robustly binds near *ppsA* and *porAB*. Additional experiments could reveal whether these genes are differentially expressed in *Har. hispanica* at other points in the growth curve, or whether these binding sites have lost their regulatory function over the course of evolution [88, 89].

The *Har. hispanica* genome encodes two GAPDH homologs belonging to the bacterial type I and archaeal type II clades (*gapI* and *gapII*, respectively). In Archaea, *gapI* homologs occur almost exclusively in saccharolytic haloarchaea and are derived from an ancient horizontal transfer of *gapI* from bacteria [34]. This acquisition, along with an amphibolic phosphoglycerate kinase (*pgk*), grants an additional ATP generation during glycolysis relative to traditional archaeal spED pathways [5]. In *Haloferax volcanii*, *gapII* and *gapI* catalyze the gluconeogenic and glycolytic reactions, respectively [34]. Our data suggest that the functional specificity of GAPDH homologs is preserved in *Har. hispanica*: *gapII* is a direct target of TrmB and its expression is strongly induced in gluconeogenic conditions. In contrast, *gapI* and cotranscribed *pgk*, are repressed in a manner consistent with TrmB-dependent regulation, though it did not pass our significance threshold (S8 FigA, S1 File). In *Har. hispanica*, TrmB-dependent activation of *gapII* is necessary for growth under gluconeogenic conditions (Fig 2 and 5), perhaps replacing regulation of *ppsA*/*pykA* as a critical metabolic control point.

Phenotype data also emphasize *gapII* as a key site of regulation: glycerol, sucrose, and fructose intermediates are predicted to enter gluconeogenesis after reactions catalyzed by GAPDHII (Fig 5C), and the addition of these carbon sources to the medium can abrogate the growth defect in Δ*trmB_Har_* background (Fig 2B). The *Har. hispanica* genome encodes a putative glucose isomerase, HAH 0464, and sucrose hydrolase, HAH 2053, which could enable the conversion of fructose and sucrose to glucose, respectively. *Har. hispanica* also encodes homologs of the bacterial-type phosphoenolpyruvate-dependent phosphotransferase system involved in fructose degradation in *Haloferax volcanii* [11, 16, 90]. These enzymes may contribute to the observed recovery of growth in the presence of sucrose and fructose, but further experiments are needed to test these predictions. In contrast, supplemental pyruvate has no effect on the growth of the deletion strain, indicating that the only way for *Har. hispanica* to generate necessary glucose from pyruvate is via the reverse EMP pathway. Interestingly, ribose has no effect on growth rate in the deletion strain, while xylose partially restores growth, suggesting *Har. hispanica* may be able to generate upper EMP intermediates from xylose but not ribose. This observation is consistent with work showing that *Har. hispanica* uses distinct enzymes to metabolize ribose and xylose and that xylose catabolism utilizes a promiscuous xylonate/gluconate dehydratase [91].

Transcriptome profiling revealed robust TrmB-dependent induction of the tryptophan biosynthesis operons, including the multi-functional fructose bisphosphate aldolase, HAH 2396. This enzyme is predicted to function in a modified shikimate synthesis pathway as well as catalyze the inter-conversion between GAP and DHAP in the EMP pathway [92]. It is evolutionarily distinct from the archaeal class I fructose bisphosphate aldolase regulated by TrmB in *Hbt. salinarum*. In *Har. hispanica*, there are no binding sites or motif occurrences in the genomic regions surrounding the tryptophan biosynthesis operons, indicating additional factors may be involved in the regulation of these promoters. A potential candidate is TrmB homolog HAH 0923, as it exhibits similar TrmB_Har_-dependent expression in response to glucose as the tryptophan operons. Unlike TrmB_Hbt_, which binds near five transcriptional regulators including itself [10], HAH 3039 and HAH 0923 are the only regulators distinguished as possible targets of TrmB in *Har. hispanica* with binding and expression data, respectively.

Due to the cryptic presence of vector sequence in the Δ*trmB_Har_* strains studied here, we primarily focus on the direct targets of TrmB_Har_ as this interaction term is better able to control for any non-specific effects of *pryF* expression and uracil prototrophy. However, we note that the expression of other metabolic pathways is altered in our data (S7 FigB). Specifically, genes involved in ribose, xylose, and arabinose catabolism are induced in Δ*trmB_Har_* exclusively when glucose is absent, a pattern consistent with indirect repression via as-yet-unknown regulators or mechanisms [91]. Expression of the operon encoding enzymes for the methylaspartate cycle is also elevated in the absence of glucose in Δ*trmB* samples, indicating that TrmB may indirectly repress other major anaplerotic pathways in *Har. hispanica*. Notably, the methylaspartate cycle in haloarchaea is correlated with the ability to synthesize the industrially-relevant biopolymer PHBV under favorable environmental conditions [93]. However, we did not observe accumulation of PHBV granules in our media conditions or any significant differential expression of the operon encoding the PHBV pathway (S8 FigB).

## Conclusion

The analyses reported here indicate that TrmB directly activates the expression of genes primarily involved in gluconeogenesis and indirectly regulates tryptophan synthesis in *Har. hispanica*, enabling cells to survive in gluconeogenic conditions. TrmB_Har_ does not directly repress the expression of glycolytic enzymes or other pathways such as cofactor biosynthesis and purine biosynthesis, or act as global transcriptional regulator similar to homologs from other archaea [10, 17]. Instead, TrmB_Har_ solely represses the expression of a putative glucose transporter when glucose is absent. Further work is needed to determine if this streamlined regulon in *Har. hispanica* is indicative of sub-functionalization and whether other TrmB homologs present may regulate some of these peripheral functions. Gluconeogenesis is an essential cellular function in haloarchaea, but it remains to be discovered whether the metabolic fate of glucose made via gluconeogenesis is conserved in metabolically distinct groups.

## Supporting information

**S1 Fig.**
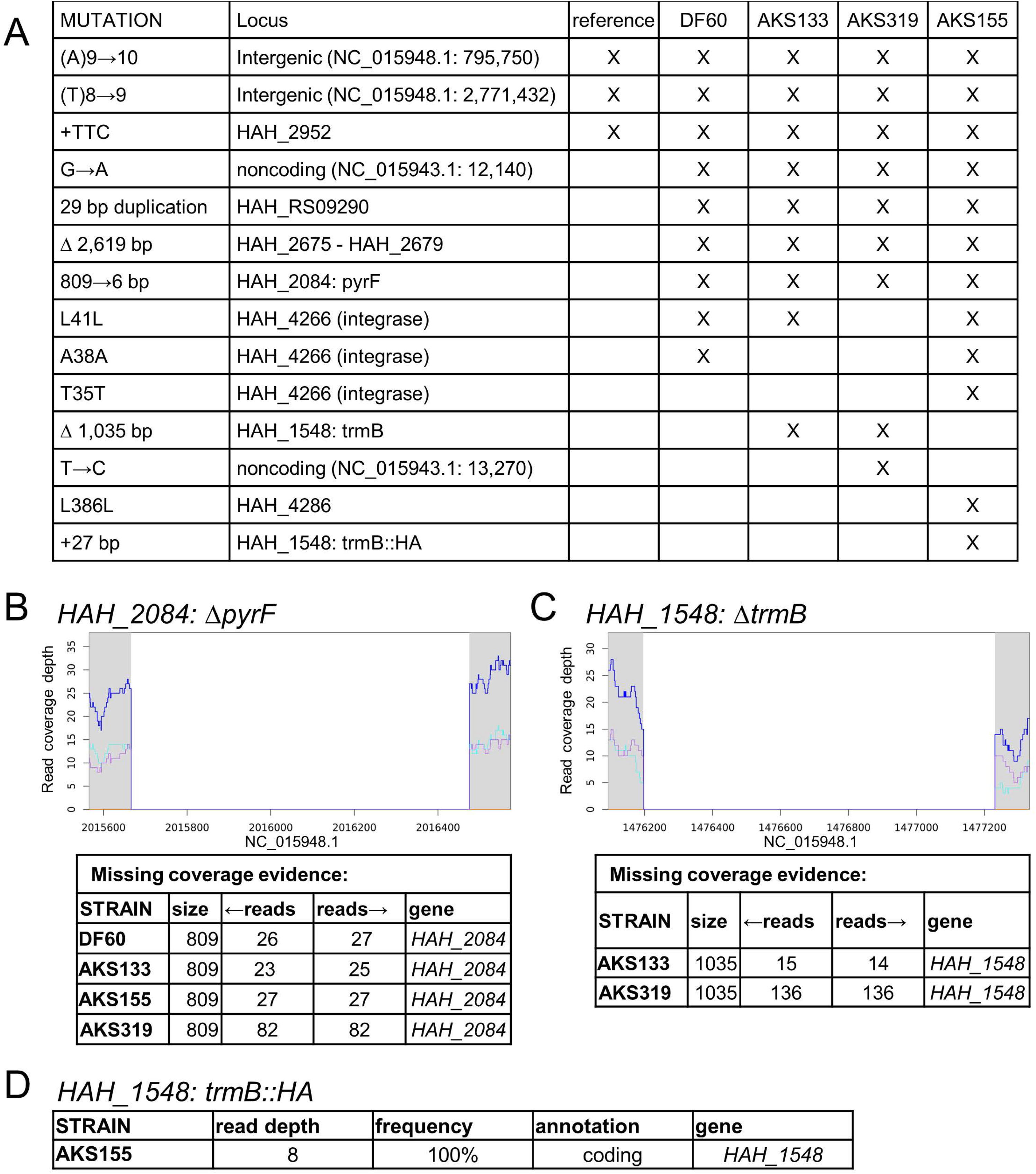
Genotype confirmation by whole-genome sequencing. A: Summary of variants identified in each strain. ”X” indicates that the mutation (rows) was detected in a given strain (columns). Strain designations are given in Supplementary Table S1. Representative coverage plots confirming chromosomal deletions for (B) *pyrF* locus for all strains and (C) *trmB_Har_* strains. X-axis provides the genome coordinates. Tables report local read depth, or sequencing coverage, for each strain. D: Confirmation of C-terminal *trmB* -hemagglutinin fusion used for immunoprecipitation experiments.

**S2 Fig.**
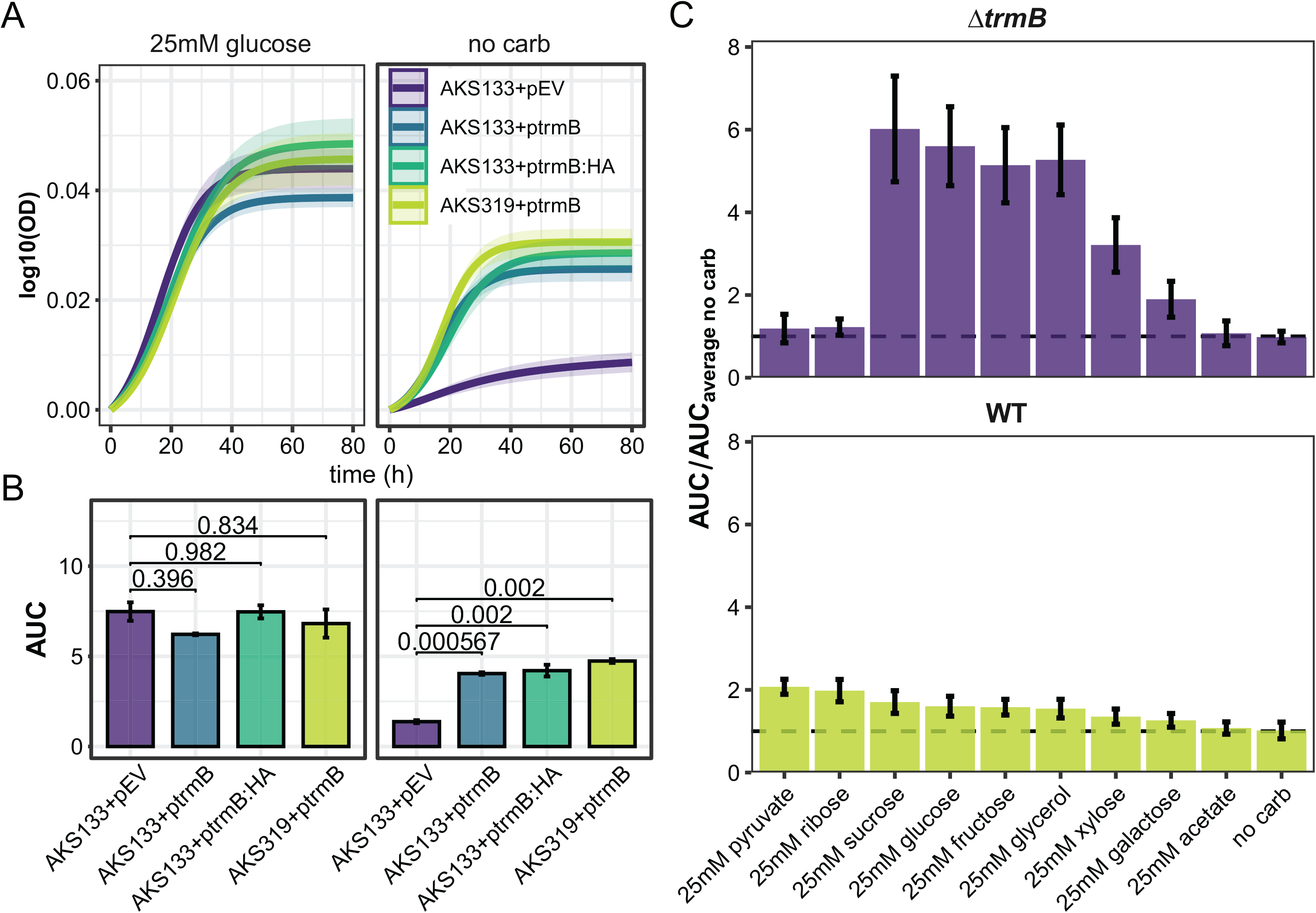
Δ*trmB* growth. A: In-trans complementation of Δ*trmB_Har_* in both AKS133 and AKS319 backgrounds. Log-transformed, fitted growth curves of complementation strains and strains harboring the empty vector (EV) grown in the presence or absence of glucose. Shaded regions depict the 95% confidence intervals. B: Area under the growth curve (AUC) of (A), with FDR-corrected significance scores. C: Parent strain and Δ*trmB_Har_* growth in each condition relative to no carbon, measured by AUC. All growth experiments were done with a minimum of 3 biological replicates, each in technical triplicate. Error bars depict the standard deviation of the mean.

**S3 Fig.**
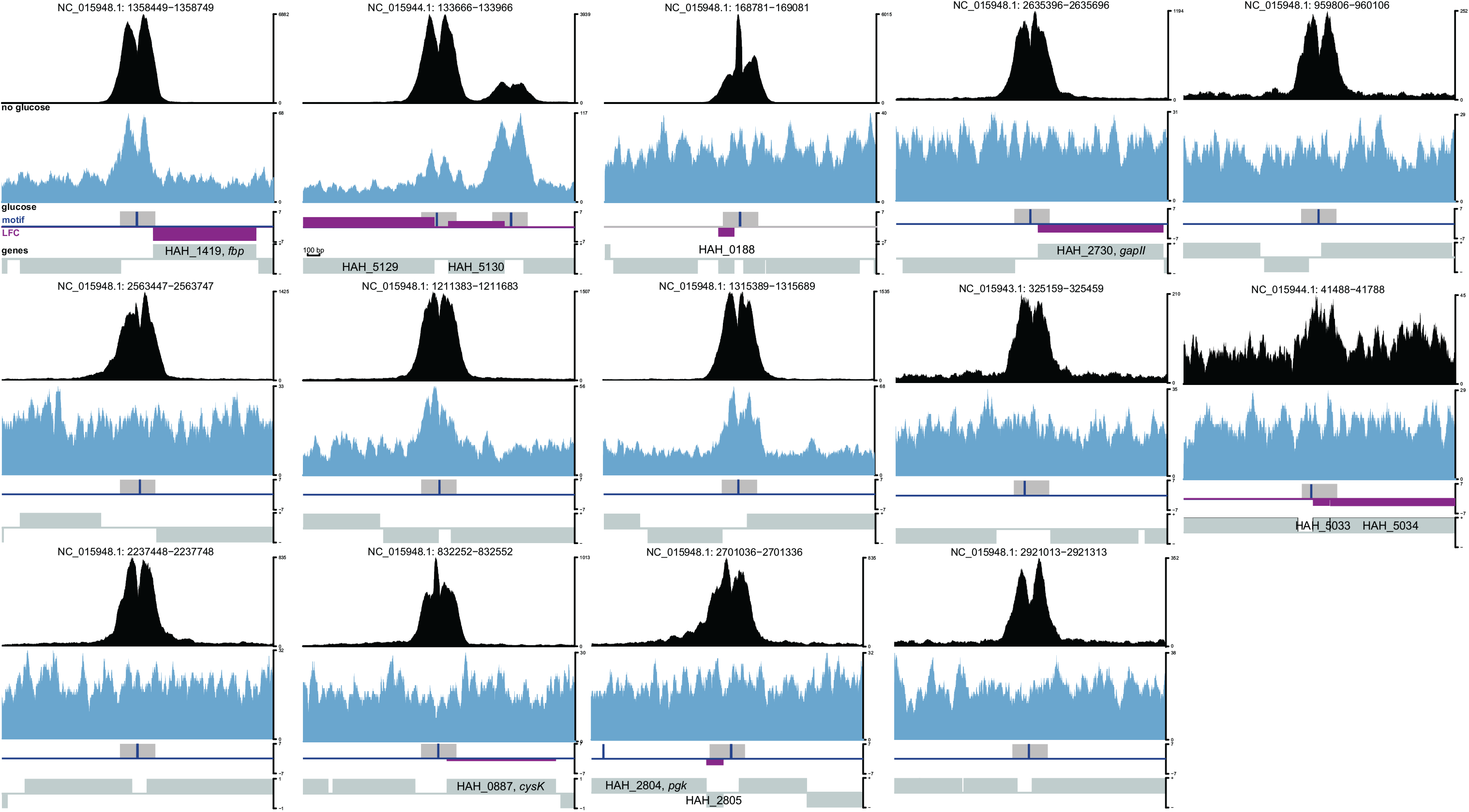
Relative position of genes, consensus peaks, and motifs for 15 regions identified by ChIP-seq. Per base coverage of a representative IP sample in the absence of glucose shown in black, an IP sample in glucose is shown in blue. The relative location of consensus peaks and motifs is shown below. Magenta bars indicate whether nearby genes were differentially expressed: magenta bars above the line represent genes up-regulated in the Δ*trmB_Har_* mutant. The height of the magenta bar represents the magnitude of change, according to the log fold change (LFC) scale bar to the right of each panel. Gene strand orientation and labels are shown in grey. Motif locations are indicated by the vertical blue line within each panel. Numbers above each panel indicate the genomic coordinates and chromosomal element of the region displayed.

**S4 Fig.**
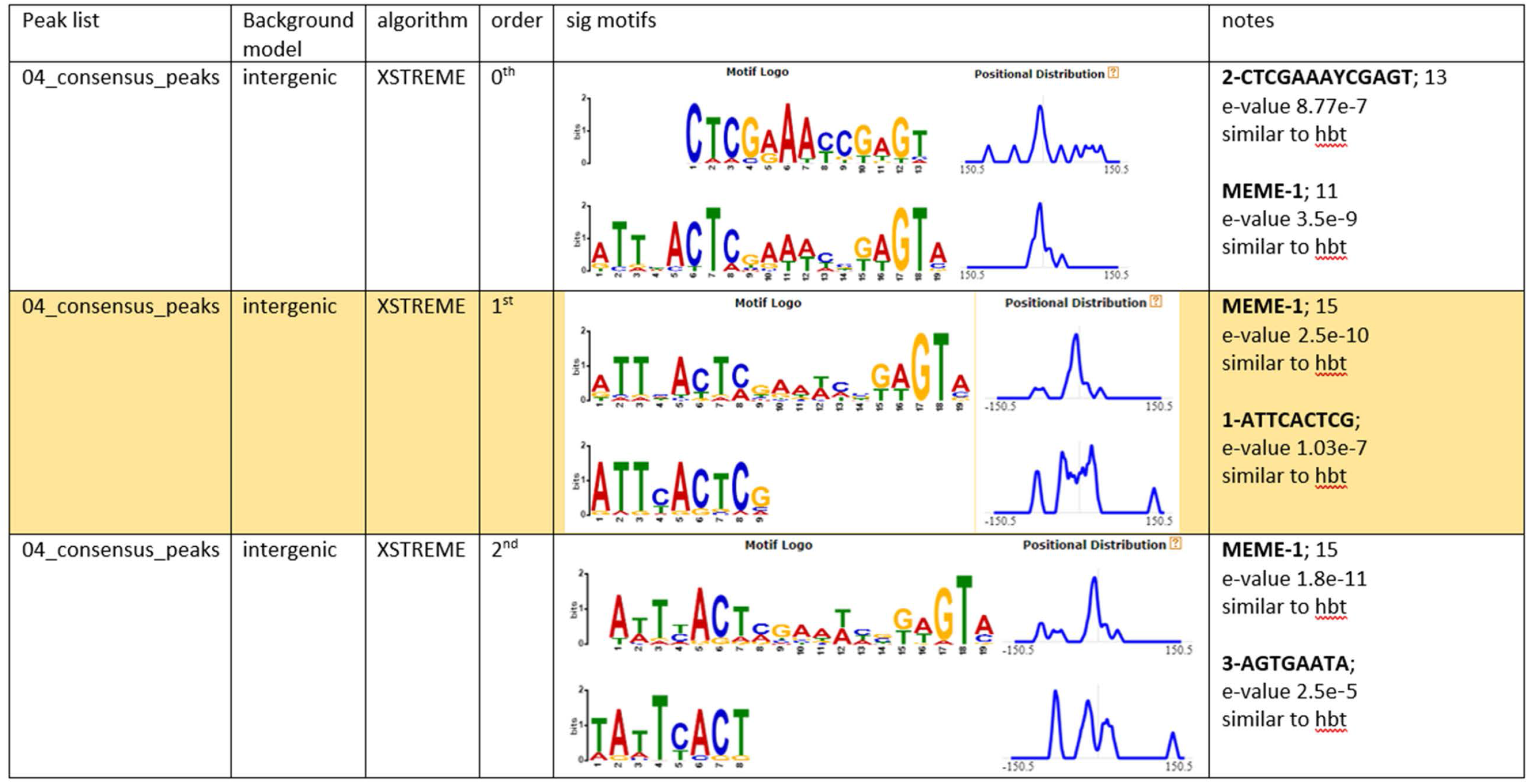
Effect of background model on discovered motifs. Sequences corresponding to peaks were extracted and submitted to XTREME as described in the methods. Yellow highlights indicate the motif reported in Fig 3.

**S5 Fig.**
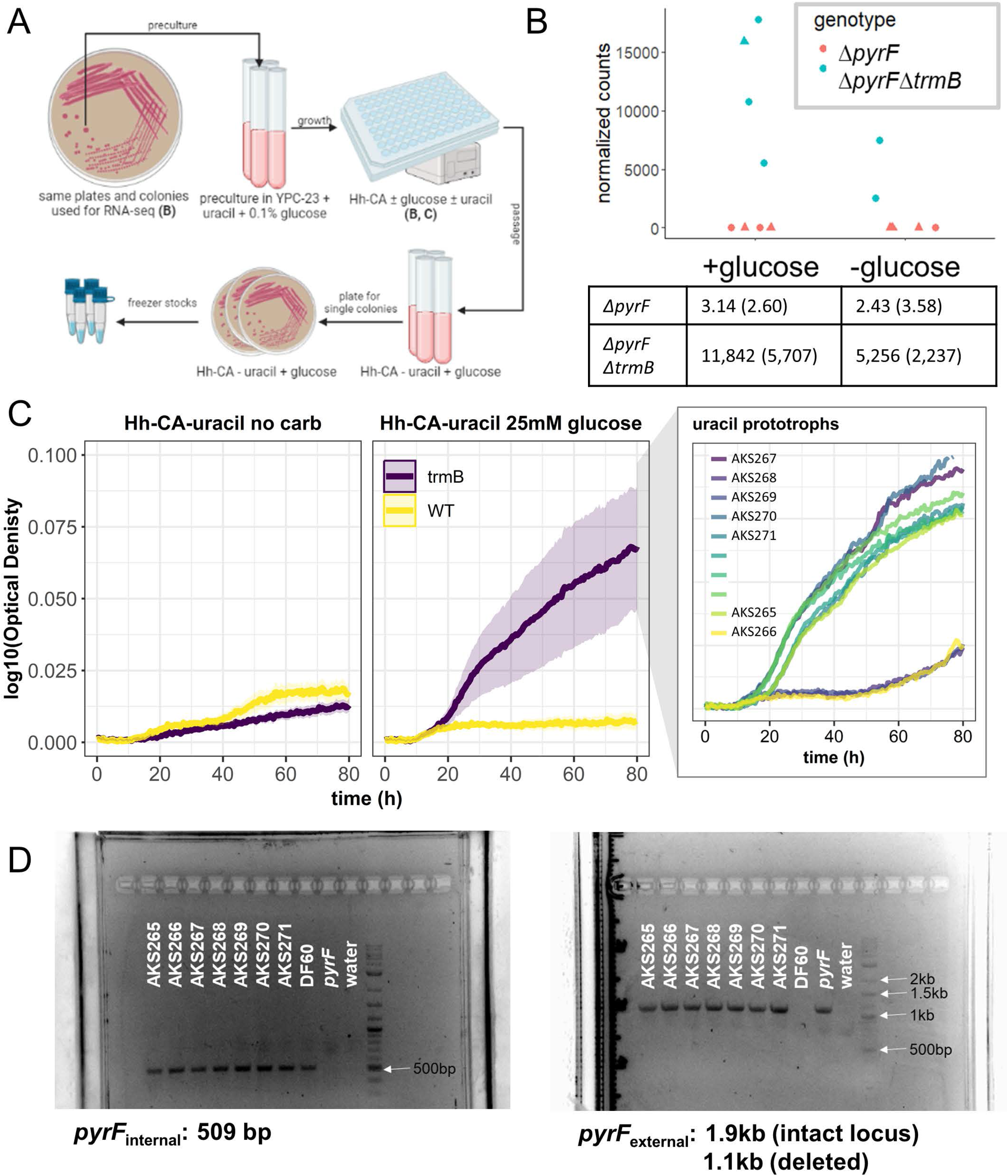
*pyrF* transcripts correspond to uracil prototrophy in AKS133. A: Diagram depicting the process of isolating uracil prototrophic strains from AKS133. B: Average number of transcripts mapping to *pyrF* in AKS133 RNA-seq samples. Standard deviation in parentheses. C: Log-transformed growth curves in the presence and absence of supplemental uracil and 5-FOA or no uracil. Shaded regions depict the 95% confidence intervals. Inset shows growth curves for individual cultures. Isolates AKS265-71 were obtained using the strategy summarized in A. D: Prototrophic isolates in C were struck from freezer stock for genomic DNA extraction. Amplification by PCR indicates *pyrF* sequence is present in the genome (left), but that the endogenous deletion is intact (right).

**S6 Fig.**
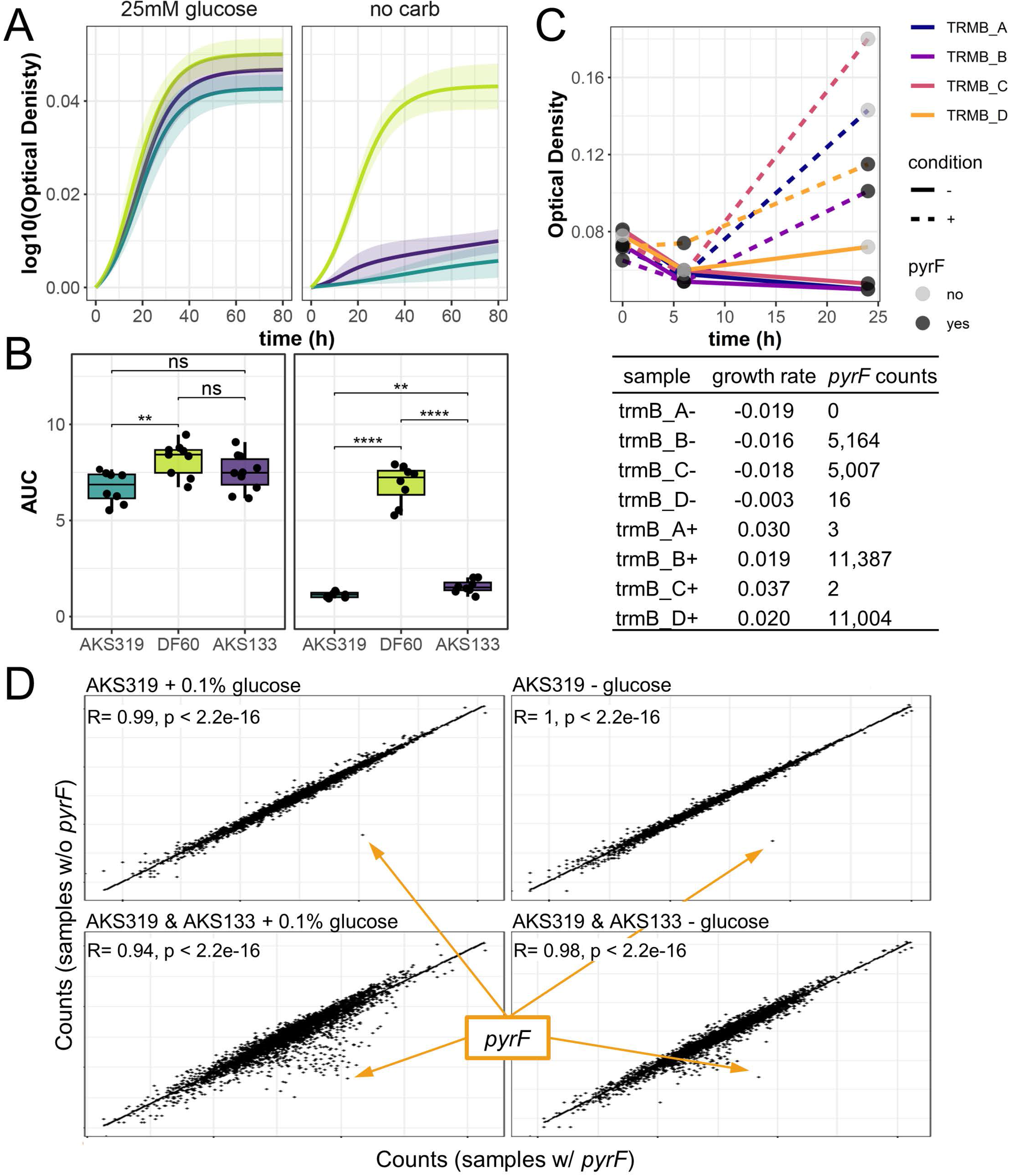
*pyrF* expression has negligible impact on Δ*trmB_Har_* fitness and transcriptome. A: Fitted, log-transformed growth curves showing that AKS319 phenocopies AKS133, and (B) that there is no significant difference between AKS133 and AKS319 in 25 mM glucose as measured by the area under the curve (AUC). Strain colors are preserved in A and B. ** *p*-value *<* 0.01; **** *<* 0.0001. C: No significant differences in the growth rate of AKS319 cultures prior to RNA extraction between replicates exhibiting *pyrF* expression and not. Optical density measurements of the cultures harvested for RNA-seq are shown, with corresponding *pyrF* counts summarized in the table below. D: Average counts per transcript are highly correlated across AKS319 samples regardless of *pyrF* expression for both -glucose (N=2) and +glucose conditions (N=2). Average counts per transcript are highly correlated across AKS319 and AKS133 regardless of *pyrF* expression for both -glucose (N=6) and +glucose conditions (N=8). Average *pryF* counts for each comparison are indicated in orange. Data are normalized relative to library size but have not been batch corrected.

**S7 Fig.**
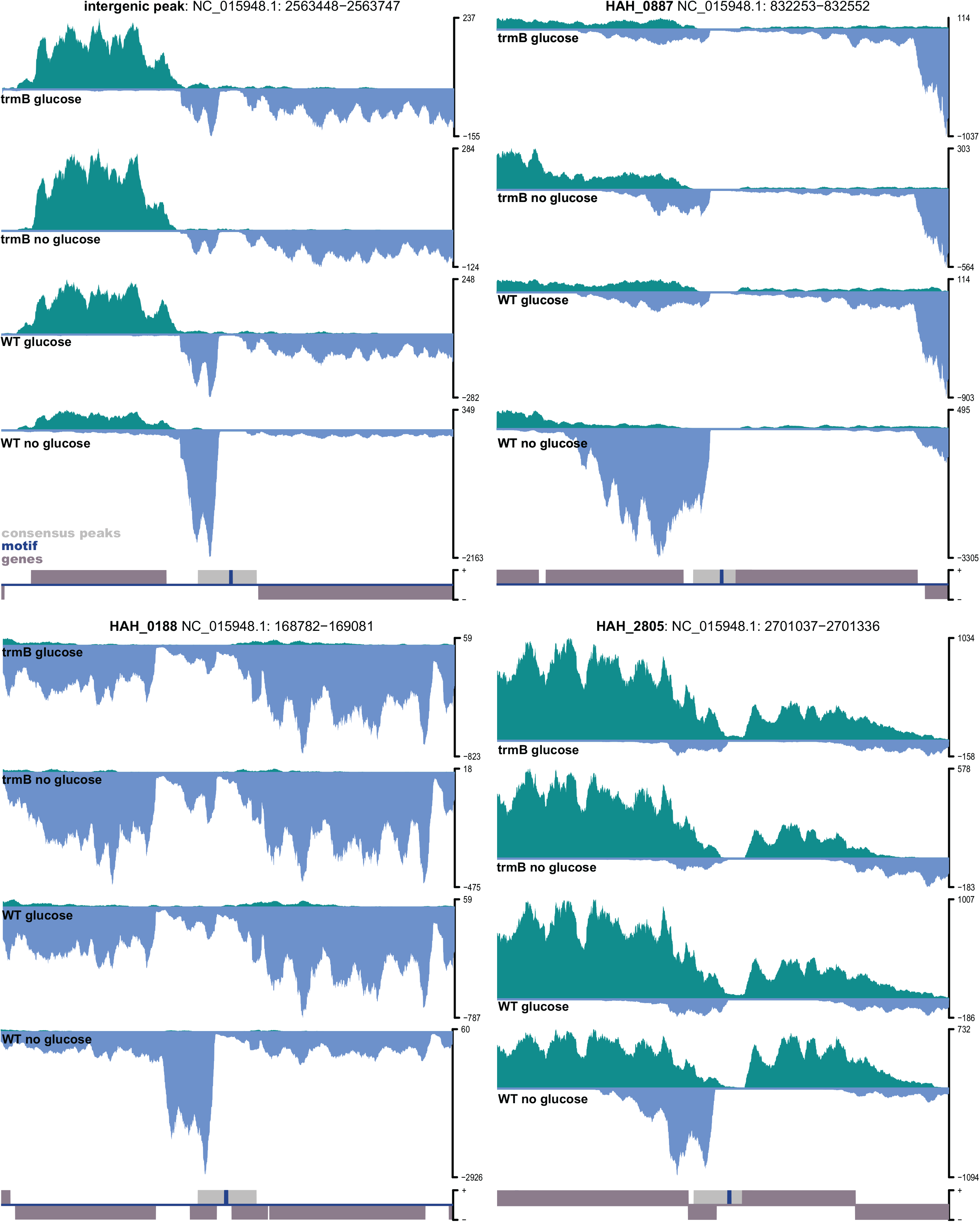
Strand-specific expression data reveal novel transcript features. Top row: Novel transcripts observed antisense to HAH 2649 (arCOG11826) and HAH 0885 (SOS response associated peptidase). Bottom row: Targets predicted to encode small proteins appear to have extended 3’ UTRs. Green tracks show the per base coverage of transcripts originating from the top strand for representative sample. Blue histograms show transcripts originating from the bottom strand.

**S8 Fig.**
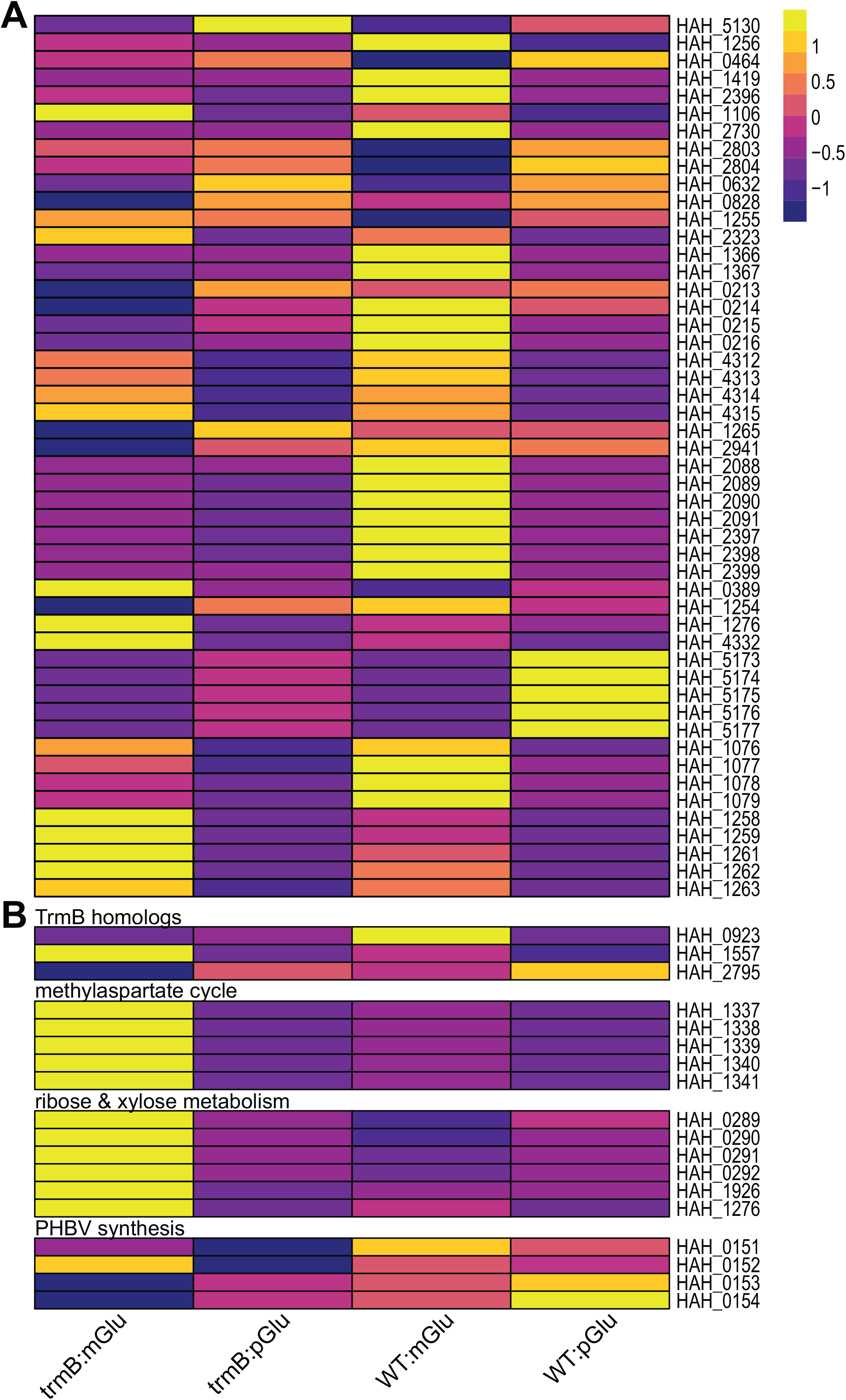
Expression of genes involved in central carbon metabolism and other catabolic pathways. A: Normalized expression pattern of genes listed in Fig 5. Yellow indicates elevated expression relative to other conditions and purple indicates reduced expression. B: Normalized expression of genes involved in characterized anaplerotic and catabolic pathways in *Har. hispanica* [13, 46, 47, 91, 93] and TrmB homologs in Table 1.

**S1 File. Supporting File 1.**

**S1 Table. Strains used in this study.**

**S2 Table. Plasmids used in this study. S3 Table. Primers used in this study.**

**S4 Table. Motif position relative to predicted transcription initiation elements for direct targets of TrmB_Har_.** Motif sequences are highlighted in purple. Motif occurrences on opposite strands were considered distinct. Darker purple color indicates motif instances on opposite strands overlap. Start codons are highlighted in grey. Putative initiation elements are bolded and underlined (TATA-box and BRE). Other haloarchaea have been reported to frequently lack identifiable TATA sequences [94]. If a promoter element could not be identified, the expected location (i.e., -26/-27 for TATA and -33/-34 for BRE) was underlined.

## Acknowledgments

*Har. hispanica* ATCC33960 and DF60 strains and the pHar plasmid were generously shared by the Xiang research group at the State Key Laboratory of Microbial Resources, Institute of Microbiology, Chinese Academy of Sciences, Beijing, China. We thank Jake Herb for his assistance in generating some of the plasmids and strains reported here, Andrew Soborowski for the list of putative transcriptional regulators in *Har. hispanica*, and Dr. Nicolas Devos and the Center for Genomic and Computational Biology at Duke University for excellent technical assistance. We acknowledge the valuable advice, comments, and feedback of current and former members of the Schmid laboratory during all stages of this project, particularly Saaz Sakrikar, Mar Martinez-Pastor, and Sungmin Hwang. Funding was provided by grants MCB-1651117, 1615685, and 1936024 from the National Science Foundation to A.K.S.

## Notes

### Competing Interest Statement

The authors have declared no competing interest.

## References

[1] Soontorngun N, Larochelle M, Drouin S, Robert F, Turcotte B. Regulation of gluconeogenesis in Saccharomyces cerevisiae is mediated by activator and repressor functions of Rds2. Mol Cell Biol. 2007;27(22):7895–7905.

[2] Morin M, Ropers D, Letisse F, Laguerre S, Portais JC, Cocaign-Bousquet M, et al. The post-transcriptional regulatory system CSR controls the balance of metabolic pools in upper glycolysis of Escherichia coli. Molecular Microbiology. 2016;100(4):686–700. https://doi.org/10.1111/mmi.13343.

[3] Wang CY, Lempp M, Farke N, Donati S, Glatter T, Link H. Metabolome and proteome analyses reveal transcriptional misregulation in glycolysis of engineered E. coli. Nat Commun. 2021;12(1):4929.

[4] Chen M, Xie T, Li H, Zhuang Y, Xia J, Nielsen J. Yeast increases glycolytic flux to support higher growth rates accompanied by decreased metabolite regulation and lower protein phosphorylation. Proceedings of the National Academy of Sciences. 2023;120(25):e2302779120. doi:10.1073/pnas.2302779120.

[5] Brasen C, Esser D, Rauch B, Siebers B. Carbohydrate metabolism in Archaea: current insights into unusual enzymes and pathways and their regulation. Microbiol Mol Biol Rev. 2014;78(1):89–175.

[6] Johnsen U, Hansen T, Schönheit P. Comparative analysis of pyruvate kinases from the hyperthermophilic archaea Archaeoglobus fulgidus, Aeropyrum pernix, and Pyrobaculum aerophilum and the hyperthermophilic bacterium Thermotoga maritima: unusual regulatory properties in hyperthermophilic archaea. J Biol Chem. 2003;278(28):25417–25427.

[7] Zhong W, Cui L, Goh BC, Cai Q, Ho P, Chionh YH, et al. Allosteric pyruvate kinase-based ”logic gate” synergistically senses energy and sugar levels in Mycobacterium tuberculosis. Nat Commun. 2017;8(1):1986.

[8] Johnsen U, Reinhardt A, Landan G, Tria FDK, Turner JM, Davies C, et al. New views on an old enzyme: allosteric regulation and evolution of archaeal pyruvate kinases. FEBS J. 2019;286(13):2471–2489.

[9] Schut GJ, Brehm SD, Datta S, Adams MW. Whole-genome DNA microarray analysis of a hyperthermophile and an archaeon: Pyrococcus furiosus grown on carbohydrates or peptides. J Bacteriol. 2003;185(13):3935–3947.

[10] Schmid AK, Reiss DJ, Pan M, Koide T, Baliga NS. A single transcription factor regulates evolutionarily diverse but functionally linked metabolic pathways in response to nutrient availability. Mol Syst Biol. 2009;5:282. doi:10.1038/msb.2009.40.

[11] Rawls KS, Yacovone SK, Maupin-Furlow JA. GlpR represses fructose and glucose metabolic enzymes at the level of transcription in the haloarchaeon *Haloferax volcanii*. J Bacteriol. 2010;192(23):6251–60. doi:10.1128/JB.00827-10.

[12] Joshua CJ, Dahl R, Benke PI, Keasling JD. Absence of diauxie during simultaneous utilization of glucose and xylose by *Sulfolobus acidocaldarius*. J Bacteriol. 2011;193(6):1293–1301.

[13] Liu H, Luo Y, Han J, Wu J, Wu Z, Feng D, et al. Proteome reference map of *Haloarcula hispanica* and comparative proteomic and transcriptomic analysis of polyhydroxyalkanoate biosynthesis under genetic and environmental perturbations. J Proteome Res. 2013;12(3):1300–15. doi:10.1021/pr300969m.

[14] López Muñoz MM, Schönheit P, Metcalf WW. Genetic, genomic, and transcriptomic studies of pyruvate metabolism in *Methanosarcina barkeri fusaro*. J Bacteriol. 2015;197(22):3592–3600.

[15] Martinez-Pastor M, Tonner PD, Darnell CL, Schmid AK. Transcriptional regulation in Archaea: from individual genes to global regulatory networks. Annu Rev Genet. 2017;51:143–170.

[16] Martin JH, Rawls KS, Chan JC, Hwang S, Martinez-Pastor M, McMillan LJ, et al. GlpR is a direct transcriptional repressor of fructose metabolic genes in *Haloferax volcanii* . J Bacteriol. 2018;200(17):e00244–18.

[17] Reichelt R, Gindner A, Thomm M, Hausner W. Genome-wide binding analysis of the transcriptional regulator TrmBL1 in Pyrococcus furiosus. BMC Genomics. 2016;17:40.

[18] Lee SJ, Surma M, Seitz S, Hausner W, Thomm M, Boos W. Characterization of the TrmB-like protein, PF0124, a TGM-recognizing global transcriptional regulator of the hyperthermophilic archaeon Pyrococcus furiosus. Mol Microbiol. 2007;65(2):305–318.

[19] Kanai T, Akerboom J, Takedomi S, van de Werken HJ, Blombach F, van der Oost J, et al. A global transcriptional regulator in Thermococcus kodakaraensis controls the expression levels of both glycolytic and gluconeogenic enzyme-encoding genes. J Biol Chem. 2007;282(46):33659–33670.

[20] Kim M, Park S, Lee SJ. Global transcriptional regulator TrmB family members in prokaryotes. J Microbiol. 2016;54(10):639–645.

[21] Gindner A, Hausner W, Thomm M. The TrmB family: a versatile group of transcriptional regulators in Archaea. Extremophiles. 2014;18(5):925–36. doi:10.1007/s00792-014-0677-2.

[22] Mondragon P, Hwang S, Kasirajan L, Oyetoro R, Nasthas A, Winters E, et al. TrmB Family Transcription Factor as a Thiol-Based Regulator of Oxidative Stress Response. mBio. 2022;13(4):e0063322.

[23] Todor H, Sharma K, Pittman AM, Schmid AK. Protein-DNA binding dynamics predict transcriptional response to nutrients in archaea. Nucleic Acids Res. 2013;41(18):8546–58. doi:10.1093/nar/gkt659.

[24] Gochnauer MB, Kushner DJ. Growth and nutrition of extremely halophilic bacteria. Can J Microbiol. 1969;15(10):1157–1165.

[25] Sumper M. Halobacterial glycoprotein biosynthesis. Biochim Biophys Acta. 1987;906(1):69–79.

[26] Severina LO, Pimenov NV, Plakunov VK. Glucose transport into the extremely halophilic archaebacteria. Arch Microbiol. 1991;155:131–136.

[27] Sonawat HM, Srivastava S, Swaminathan S, Govil G. Glycolysis and Entner-Doudoroff pathways in Halobacterium halobium: some new observations based on 13C NMR spectroscopy. Biochem Biophys Res Commun. 1990;173(1):358–362.

[28] Gruber C, Legat A, Pfaffenhuemer M, Radax C, Weidler G, Busse HJ, et al. Halobacterium noricense sp. nov., an archaeal isolate from a bore core of an alpine Permian salt deposit, classification of Halobacterium sp. NRC-1 as a strain of H. salinarum and emended description of H. salinarum. Extremophiles. 2004;8(6):431–439.

[29] Todor H, Dulmage K, Gillum N, Bain JR, Muehlbauer MJ, Schmid AK. A transcription factor links growth rate and metabolism in the hypersaline adapted archaeon Halobacterium salinarum. Mol Microbiol. 2014;93(6):1172–82. doi:10.1111/mmi.12726.

[30] Todor H, Gooding J, Ilkayeva OR, Schmid AK. Dynamic Metabolite Profiling in an Archaeon Connects Transcriptional Regulation to Metabolic Consequences. PLoS One. 2015;10(8):e0135693. doi:10.1371/journal.pone.0135693.

[31] Tomlinson GA, Koch TK, Hochstein LI. The metabolism of carbohydrates by extremely halophilic bacteria: glucose metabolism via a modified Entner-Doudoroff pathway. Can J Microbiol. 1974;20(8):1085–1091.

[32] Johnsen U, Selig M, Xavier KB, Santos H, Schönheit P. Different glycolytic pathways for glucose and fructose in the halophilic archaeon Halococcus saccharolyticus. Arch Microbiol. 2001;175(1):52–61.

[33] Sutter JM, Tästensen JB, Johnsen U, Soppa J, Schönheit P. Key Enzymes of the Semiphosphorylative Entner-Doudoroff Pathway in the Haloarchaeon Haloferax volcanii: Characterization of Glucose Dehydrogenase, Gluconate Dehydratase, and 2-Keto-3-Deoxy-6-Phosphogluconate Aldolase. J Bacteriol. 2016;198(16):2251–2262.

[34] Tästensen JB, Schönheit P. Two distinct glyceraldehyde-3-phosphate dehydrogenases in glycolysis and gluconeogenesis in the archaeon Haloferax volcanii. FEBS Lett. 2018;592(9):1524–1534.

[35] Gonzalez O, Gronau S, Pfeiffer F, Mendoza E, Zimmer R, Oesterhelt D. Systems analysis of bioenergetics and growth of the extreme halophile Halobacterium salinarum. PLoS Comput Biol. 2009;5(4):e1000332. doi:10.1371/journal.pcbi.1000332.

[36] Anderson I, Scheuner C, Göker M, Mavromatis K, Hooper SD, Porat I, et al. Novel insights into the diversity of catabolic metabolism from ten haloarchaeal genomes. PLoS One. 2011;6(5):e20237.

[37] Feng J, Liu B, Zhang Z, Ren Y, Li Y, Gan F, et al. The complete genome sequence of Natrinema sp. J7-2, a haloarchaeon capable of growth on synthetic media without amino acid supplements. PLoS One. 2012;7(7):e41621.

[38] Rawal N, Kelkar SM, Altekar W. Alternative routes of carbohydrate metabolism in halophilic archaebacteria. Indian J Biochem Biophys. 1988;25(6):674–686.

[39] Amoozegar MA, Siroosi M, Atashgahi S, Smidt H, Ventosa A. Systematics of haloarchaea and biotechnological potential of their hydrolytic enzymes. Microbiology (Reading). 2017;163(5):623–645.

[40] Mitra R, Xu T, Xiang H, Han J. Current developments on polyhydroxyalkanoates synthesis by using halophiles as a promising cell factory. Microb Cell Fact. 2020;19(1):86.

[41] Torreblanca M, Rodriguez-Valera F, Juez G, Ventosa A, Kamekura M, Kates M. Classification of Non-alkaliphilic Halobacteria Based on Numerical Taxonomy and Polar Lipid Composition, and Description of Haloarcula gen. nov. and Haloferax gen. nov. Systematic and Applied Microbiology. 1986;8(1):89–99. https://doi.org/10.1016/S0723-2020(86)80155-2.

[42] Juez G, Rodriguez-Valera F, Ventosa A, Kushner DJ. Haloarcula hispanica spec. nov. and Haloferax gibbonsii spec, nov., two new species of extremely halophilic archaebacteria. Systematic and Applied Microbiology. 1986;8(1-2):75–79.

[43] Liu H, Wu Z, Li M, Zhang F, Zheng H, Han J, et al. Complete genome sequence of Haloarcula hispanica, a Model Haloarchaeon for studying genetics, metabolism, and virus-host interaction. J Bacteriol. 2011;193(21):6086–7. doi:10.1128/JB.05953-11.

[44] Liu H, Han J, Liu X, Zhou J, Xiang H. Development of pyrF-based gene knockout systems for genome-wide manipulation of the archaea Haloferax mediterranei and Haloarcula hispanica. J Genet Genomics. 2011;38(6):261–9. doi:10.1016/j.jgg.2011.05.003.

[45] Han J, Lu Q, Zhou L, Zhou J, Xiang H. Molecular characterization of the phaECHm genes, required for biosynthesis of poly(3-hydroxybutyrate) in the extremely halophilic archaeon Haloarcula marismortui. Appl Environ Microbiol. 2007;73(19):6058–65. doi:10.1128/AEM.00953-07.

[46] Han J, Lu Q, Zhou L, Liu H, Xiang H. Identification of the polyhydroxyalkanoate (PHA)-specific acetoacetyl coenzyme A reductase among multiple FabG paralogs in *Haloarcula hispanica* and reconstruction of the PHA biosynthetic pathway in *Haloferax volcanii*. Appl Environ Microbiol. 2009;75(19):6168–75. doi:10.1128/AEM.00938-09.

[47] Han J, Hou J, Liu H, Cai S, Feng B, Zhou J, et al. Wide distribution among halophilic archaea of a novel polyhydroxyalkanoate synthase subtype with homology to bacterial type III synthases. Appl Environ Microbiol. 2010;76(23):7811–7819.

[48] Allers T, Ngo HP, Mevarech M, Lloyd RG. Development of additional selectable markers for the halophilic archaeon Haloferax volcanii based on the leuB and trpA genes. Appl Environ Microbiol. 2004;70(2):943–953.

[49] Gibson DG. Enzymatic assembly of overlapping DNA fragments. Meth Enzymol. 2011;498:349–361.

[50] Cai S, Cai L, Liu H, Liu X, Han J, Zhou J, et al. Identification of the haloarchaeal phasin (PhaP) that functions in polyhydroxyalkanoate accumulation and granule formation in Haloferax mediterranei. Appl Environ Microbiol. 2012;78(6):1946–52. doi:10.1128/AEM.07114-11.

51. Dyall-Smith M. The Halohandbook: Protocols for haloarchaeal genetics; 2009. Available from: https://haloarchaea.com/wp-content/uploads/2018/10/Halohandbook_2009_v7.3mds.pdf.

[52] Deatherage DE, Barrick JE. Identification of mutations in laboratory-evolved microbes from next-generation sequencing data using breseq. Methods Mol Biol. 2014;1151:165–188.

[53] Finn RD, Mistry J, Schuster-Bockler B, Griffiths-Jones S, Hollich V, Lassmann T, et al. Pfam: clans, web tools and services. Nucleic Acids Res. 2006;34(Database issue):D247–51.

[54] Madeira F, Pearce M, Tivey ARN, Basutkar P, Lee J, Edbali O, et al. Search and sequence analysis tools services from EMBL-EBI in 2022. Nucleic Acids Res. 2022;50(W1):W276–279.

[55] Sprouffske K, Wagner A. Growthcurver: an R package for obtaining interpretable metrics from microbial growth curves. BMC Bioinformatics. 2016;17:172.

[56] Das S, Noe JC, Paik S, Kitten T. An improved arbitrary primed PCR method for rapid characterization of transposon insertion sites. J Microbiol Methods. 2005;63(1):89–94.

[57] Wilbanks EG, Larsen DJ, Neches RY, Yao AI, Wu CY, Kjolby RA, et al. A workflow for genome-wide mapping of archaeal transcription factors with ChIP-seq. Nucleic Acids Res. 2012;40(10):e74. doi:10.1093/nar/gks063.

[58] Langmead B, Salzberg SL. Fast gapped-read alignment with Bowtie 2. Nat Methods. 2012;9(4):357–9. doi:10.1038/nmeth.1923.

[59] Li H, Handsaker B, Wysoker A, Fennell T, Ruan J, Homer N, et al. The Sequence Alignment/Map format and SAMtools. Bioinformatics. 2009;25(16):2078–2079.

[60] Carroll TS, Liang Z, Salama R, Stark R, de Santiago I. Impact of artifact removal on ChIP quality metrics in ChIP-seq and ChIP-exo data. Front Genet. 2014;5:75.

[61] Ross-Innes CS, Stark R, Teschendorff AE, Holmes KA, Ali HR, Dunning MJ, et al. Differential oestrogen receptor binding is associated with clinical outcome in breast cancer. Nature. 2012;481(7381):389–393.

[62] Ou J, Zhu LJ. trackViewer: a Bioconductor package for interactive and integrative visualization of multi-omics data. Nat Methods. 2019;16(6):453–454.

[63] Lawrence M, Huber W, s H, Aboyoun P, Carlson M, Gentleman R, et al. Software for computing and annotating genomic ranges. PLoS Comput Biol. 2013;9(8):e1003118.

[64] Grünberger F, Reichelt R, Waege I, Ned V, Bronner K, Kaljanac M, et al. CopR, a Global Regulator of Transcription to Maintain Copper Homeostasis in Pyrococcus furiosus. Front Microbiol. 2020;11:613532.

[65] Pastor MM, Sakrikar S, Rodriguez DN, Schmid AK. Comparative Analysis of rRNA Removal Methods for RNA-Seq Differential Expression in Halophilic Archaea. Biomolecules. 2022;12(5).

[66] Liao Y, Smyth GK, Shi W. featureCounts: an efficient general purpose program for assigning sequence reads to genomic features. Bioinformatics. 2014;30(7):923–930.

[67] Love MI, Huber W, Anders S. Moderated estimation of fold change and dispersion for RNA-seq data with DESeq2. Genome Biol. 2014;15(12):550. doi:10.1186/s13059-014-0550-8.

[68] Sakrikar S, Schmid AK. An archaeal histone-like protein regulates gene expression in response to salt stress. Nucleic Acids Res. 2021;49(22):12732–12743.

[69] Huerta-Cepas J, Forslund K, Coelho LP, Szklarczyk D, Jensen LJ, von Mering C, et al. Fast genome-wide functional annotation through orthology assignment by eggNOG-mapper. Mol Biol Evol. 2017;34(8):2115–2122.

70. Grant CE, Bailey TL. XSTREME: Comprehensive motif analysis of biological sequence datasets. bioRxiv. 2021;doi:10.1101/2021.09.02.458722.

[71] Mistry J, Chuguransky S, Williams L, Qureshi M, Salazar GA, Sonnhammer ELL, et al. Pfam: The protein families database in 2021. Nucleic Acids Res. 2021;49(D1):D412–D419.

[72] Krug M, Lee SJ, Diederichs K, Boos W, Welte W. Crystal structure of the sugar binding domain of the archaeal transcriptional regulator TrmB. J Biol Chem. 2006;281(16):10976–10982.

[73] Krug M, Lee SJ, Boos W, Diederichs K, Welte W. The three-dimensional structure of TrmB, a transcriptional regulator of dual function in the hyperthermophilic archaeon Pyrococcus furiosus in complex with sucrose. Protein Sci. 2013;22(6):800–808.

[74] Koide T, Reiss DJ, Bare JC, Pang WL, Facciotti MT, Schmid AK, et al. Prevalence of transcription promoters within archaeal operons and coding sequences. Mol Syst Biol. 2009;5:285. doi:10.1038/msb.2009.42.

[75] Breuert S, Allers T, Spohn G, Soppa J. Regulated polyploidy in halophilic archaea. PLoS One. 2006;1:e92.

76. Schiller H, Kouassi J, Hong Y, Rados T, Kwak J, DiLucido A, et al. Identification and characterization of structural and regulatory cell-shape determinants in Haloferax volcanii. bioRxiv. 2023;doi:10.1101/2023.03.05.531186.

[77] van de Werken HJ, Verhees CH, Akerboom J, de Vos WM, van der Oost J. Identification of a glycolytic regulon in the archaea Pyrococcus and Thermococcus. FEMS Microbiol Lett. 2006;260(1):69–76. doi:10.1111/j.1574-6968.2006.00292.x.

[78] Gupta S, Stamatoyannopoulos JA, Bailey TL, Noble WS. Quantifying similarity between motifs. Genome Biol. 2007;8(2):R24.

[79] Bell SD, Cairns SS, Robson RL, Jackson SP. Transcriptional regulation of an archaeal operon in vivo and in vitro. Mol Cell. 1999;4(6):971–982.

[80] Hain J, Reiter WD, Hudepohl U, Zillig W. Elements of an archaeal promoter defined by mutational analysis. Nucleic Acids Res. 1992;20(20):5423–5428.

[81] Paya G, Bautista V, Camacho M, Castejon-Fernandez N, Alcaraz LA, Bonete MJ, et al. Small RNAs of *Haloferax mediterranei* : identification and potential involvement in nitrogen metabolism. Genes (Basel). 2018;9(2).

[82] Gelsinger DR, DiRuggiero J. Transcriptional Landscape and Regulatory Roles of Small Noncoding RNAs in the Oxidative Stress Response of the Haloarchaeon *Haloferax volcanii* . J Bacteriol. 2018;200(9):e00779–17. doi:10.1128/JB.00779-17.

[83] Kliemt J, Jaschinski K, Soppa J. A Haloarchaeal Small Regulatory RNA (sRNA) Is Essential for Rapid Adaptation to Phosphate Starvation Conditions. Front Microbiol. 2019;10:1219.

[84] Gelsinger DR, Reddy R, Whittington K, Debic S, DiRuggiero J. Post-transcriptional regulation of redox homeostasis by the small RNA SHOxi in haloarchaea. RNA Biol. 2021;18(11):1867–1881.

[85] Prasse D, Förstner KU, Jäger D, Backofen R, Schmitz RA. 1. RNA Biol. 2017;14(11):1544–1558.

[86] Lee SJ, Surma M, Seitz S, Hausner W, Thomm M, Boos W. Differential signal transduction via TrmB, a sugar sensing transcriptional repressor of Pyrococcus furiosus. Mol Microbiol. 2007;64(6):1499–1505.

[87] Kuprat T, Ortjohann M, Johnsen U, Schönheit P. Glucose Metabolism and Acetate Switch in Archaea: the Enzymes in Haloferax volcanii. J Bacteriol. 2021;203(8).

[88] Fitzgerald DM, Smith C, Lapierre P, Wade JT. The evolutionary impact of intragenic FliA promoters in proteobacteria. Mol Microbiol. 2018;108(4):361–378.

[89] Nguyen-Duc T, van Oeffelen L, Song N, Hassanzadeh-Ghassabeh G, Muyldermans S, Charlier D, et al. The genome-wide binding profile of the Sulfolobus solfataricus transcription factor Ss-LrpB shows binding events beyond direct transcription regulation. BMC Genomics. 2013;14(1):828.

[90] Pickl A, Johnsen U, Schönheit P. Fructose degradation in the haloarchaeon Haloferax volcanii involves a bacterial type phosphoenolpyruvate-dependent phosphotransferase system, fructose-1-phosphate kinase, and class II fructose-1,6-bisphosphate aldolase. J Bacteriol. 2012;194(12):3088–3097.

[91] Johnsen U, Sutter JM, Reinhardt A, Pickl A, Wang R, Xiang H, et al. d-Ribose Catabolism in Archaea: Discovery of a Novel Oxidative Pathway in Haloarcula Species. J Bacteriol. 2020;202(3).

[92] Hochuli M, Patzelt H, Oesterhelt D, Wüthrich K, Szyperski T. Amino acid biosynthesis in the halophilic archaeon Haloarcula hispanica. J Bacteriol. 1999;181(10):3226–3237.

[93] Borjian F, Han J, Hou J, Xiang H, Berg IA. The methylaspartate cycle in haloarchaea and its possible role in carbon metabolism. ISME J. 2016;10(3):546–557.

[94] Martinez GS, Sarkar S, Kumar A, rez Rueda E, de Avila E Silva S. Characterization of promoters in archaeal genomes based on DNA structural parameters. Microbiologyopen. 2021;10(5):e1230.

