## supplemental tables 1-3 for "A conserved transcription factor controls gluconeogenesis via distinct targets in hypersaline-adapted archaea with diverse metabolic capabilities"

**Supplementary Table 1.** Primers used in this study.

| Primer name | Sequence (5’ 🡪 3’) | Purpose |
| --- | --- | --- |
| Hh_1548_up_F | TCCGCTAAGGTACCTCTAGAAGAAGCTTGGGGAACCGACGGCTACTATGC | trmB deletion, locus amplification for integrant tagging |
| Hh_1548_up_R | CTACAGCCGACAGTGTCAGTTCTCACTAGACATAGTGGAGCGTTTGCG | trmB deletion |
| Hh_1548_down_F | CGCAAACGCTCCACTATGTCTAGTGAGAACTGACACTGTCGGCTGTAG | trmB deletion |
| Hh_1548_down_R | AGGGCCCCTGCAGGTCGACTCTAGAGGATCCTCGCTCGCAGTGACCGC | trmB deletion, locus amplification for integrant tagging |
| Hh_trmBHA_F2 | gccggattatgcgTGACACTGTCGGCTGTAG | HA-tag insertion |
| Hh_trmBHA_R2 | acatcatacggataGTTCTCGGGCGGTTCGTC | HA-tag insertion |
| MevR_BamHI_F | AAAGGATCCGGGTGTGTACCTCCGCGTTCGTC | Mev^R^ template for pAKS83 |
| MevR_SmaI_R | AAACCCGGGTTACCGACCGAGTTCGGCGTGGG | Mev^R^ template for pAKS83 |
| Hh_PtrmB_EcoRI_F | AAAgaattcCCGCTATCTCATCGCTGAAC | trmB and trmB-HA complementation plasmids |
| Hh_trmB_HindIII_R | TTTaagcttTCAGTTCTCGGGCGGTTCGTC | trmB complementation plasmid |
| HA_HindIII_R | TTTaagcttTCACGCATAATCCGGCACATCA | trmB-HA complementation plasmids |
| Hh_trmB_F | ATGTCTAGTGACGACTTGGAAGC | Sanger sequencing, plasmid verification |
| pHAR_MCS_1 | GACAGTCCGCGAAACAGCTC | Sanger sequencing, plasmid verification |
| pHAR_F | ACGACTCCGGTGACGCGTTCTTCA | Sanger sequencing, plasmid verification |
| pHAR_R | CATGATTACGCCAGATATCAAATT | Sanger sequencing, plasmid verification |
| Hh_idr1_F | CTTCGGATCCAGGACGAGCAGTTTGACAGC | gDNA contamination PCR |
| Hh_idr1_R | CTGTGGATCCTTCGATGGTATCACCGACCT | gDNA contamination PCR |
| Hh_pyrFint_R3 | AACTGATTCAGTCGCTGTTTG | Amplify 509bp region of pyrF CDS |
| Hh_pyrFint_F4 | ATTCTCGACGACGAGGAAGG | Amplify 509bp region of pyrF CDS |
| Hh_pyrF_PCR_F | CGACTCGGCTCGGCAATA | Amplify over the endogenous pyrF locus |
| Hh_pyrF_PCR_R | CCAGCATTCCGAGTATCCA | Amplify over the endogenous pyrF locus |

**Supplementary Table 2.** Plasmids used in this study

| Name | Description | Species | Reference |
| --- | --- | --- | --- |
| pHar | integration vector; Amp^R^, *pyrF* | *Haloarcula hispanica* | Lui 2011. 10.1016/j.jgg.2011.05.003 |
| pWL502 | derivative of pWL102; replaced Mev^R^ with *pyrF* marker | *Haloferax mediterranei* | Cai et al, 2012. doi:10.1128/aem.07114-11 |
| pNBKO7 | Template for pAKS83 Mev^R^ cassette | *Halobacterium salinarum* | Willebanks 2012. 10.1093/nar/gks063 |
| pAKS139 | pHar with *trmB* flanking regions for deletion | *Haloarcula hispanica* | This study |
| pAKS76 | pHar with trmB locus at BamHI site | *Haloarcula hispanica* | This study |
| pAKS87 | pHar with trmB-HA fusion at BamHI site | *Haloarcula hispanica* | This study |
| pAKS83 | pWL502 based self-replicating plasmid with Mev^R^ gene for selection | *Haloarcula hispanica* | This study |
| pAKS95 | pAKS83-based construct carrying *trmB* under its native promoter at the EcoRI and HindIII sites. | *Haloarcula hispanica* | This study |
| pAKS192 | pAKS83-based construct carrying *trmB-HA* translational fusion under its native promoter at the EcoRI and HindIII sites. | *Haloarcula hispanica* | This study |

**Supplementary Table 3.** Strains used in this study

| Name | Species | Genotype | Purpose | Reference |
| --- | --- | --- | --- | --- |
| DF60 | *Haloarcula hispanica* ATCC33960 | Δ*pyrF* | growth assays, whole genome sequencing, ChIP-seq, RNA-seq | (Liu *et al.*, 2011) |
| AKS133 | *Haloarcula hispanica* | Δ*pyrF* Δ*trmB* | Growth assays, whole genome sequencing, RNA-seq | This study |
| AKS155 | *Haloarcula hispanica* | Δ*pyrF trmB-HA* | ChIP-seq, whole genome sequencing | This study |
| AKS248 | *Haloarcula hispanica* | AKS133 *+* pAKS192 | TrmB-HA exogenous expression for complementation growth assays. | This study |
| AKS319 | *Haloarcula hispanica* | Δ*pyrF* Δ*trmB* | whole genome sequencing, RNA-seq | This study |
| AKS336 | *Haloarcula hispanica* | AKS319 *+* pAKS95 | TrmB exogenous expression for complementation growth assays. | This study |
| AKS265 | *Haloarcula hispanica* | Δ*pyrF* Δ*trmB* | AKS133 uracil prototroph isolated from well 53 | This study |
| AKS266 | *Haloarcula hispanica* | Δ*pyrF* Δ*trmB* | AKS133 uracil prototroph isolated from well 68 | This study |
| AKS267 | *Haloarcula hispanica* | Δ*pyrF* Δ*trmB* | AKS133 uracil prototroph isolated from well 125 | This study |
| AKS268 | *Haloarcula hispanica* | Δ*pyrF* Δ*trmB* | AKS133 uracil prototroph isolated from well 138 | This study |
| AKS269 | *Haloarcula hispanica* | Δ*pyrF* Δ*trmB* | AKS133 uracil prototroph isolated from well 143 | This study |
| AKS270 | *Haloarcula hispanica* | Δ*pyrF* Δ*trmB* | AKS133 uracil prototroph isolated from well 148 | This study |
| AKS271 | *Haloarcula hispanica* | Δ*pyrF* Δ*trmB* | AKS133 uracil prototroph isolated from well 158 | This study |

**Supplementary Table 4.** ChIP-Seq samples

| Sample name | Strain ID | Plate age | OD600 |
| --- | --- | --- | --- |
| A11_AH12_HH210_c3_+0.1glu_IP | AKS155 | 8 days | 0.37 |
| A8_AH4_HH37_c2_-glu_IP | DF60 | 8 days | 0.35 |
| B12_AH15_HH210_c5_-glu_WCE | AKS155 | 8 days | 0.29 |
| B4_AH10_HH210_c3_-glu_IP | AKS155 | 8 days | 0.32 |
| B5_AH13_HH210_c4_-glu_WCE | AKS155 | 8 days | 0.32 |
| C5_AH6_HH210_c2_-glu_IP | AKS155 | 8 days | 0.33 |
| D12_AH9_HH210_c3_-glu_WCE | AKS155 | 8 days | 0.32 |
| D2_AH8_HH210_c2_+0.1glu_IP | AKS155 | 8 days | 0.39 |
| D3_AH5_HH210_c2_-glu_WCE | AKS155 | 8 days | 0.33 |
| E10_AH16_HH210_c5_-glu_IP | AKS155 | 8 days | 0.29 |
| E7_AH14_HH210_c4_-glu_IP | AKS155 | 8 days | 0.32 |
| E9_AH11_HH210_c3_+0.1glu_WCE | AKS155 | 8 days | 0.37 |
| G5_AH1_HH37_c1_-glu_WCE | DF60 | 8 days | 0.36 |
| H10_AH3_HH37_c2_-glu_WCE | DF60 | 8 days | 0.35 |
| H3_AH2_HH37_c1_-glu_IP | DF60 | 8 days | 0.36 |
| H6_AH7_HH210_c2_+0.1glu_WCE | AKS155 | 8 days | 0.39 |
