## Supplementary figures and images for "A conserved transcription factor controls gluconeogenesis via distinct targets in hypersaline-adapted archaea with diverse metabolic capabilities"

### supplemental table 4

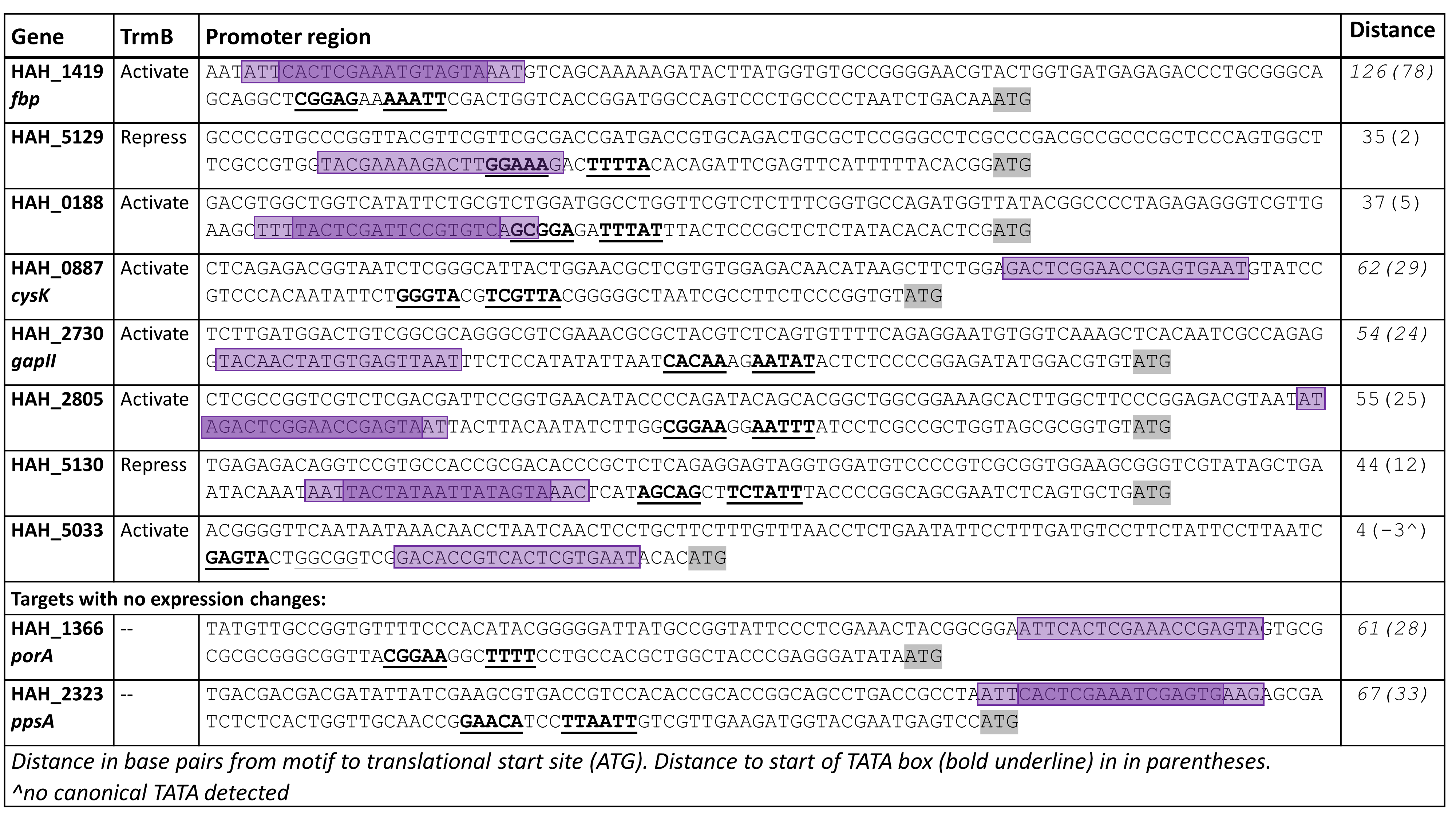
